# Macrophages employ quorum licensing to regulate collective activation

**DOI:** 10.1101/610592

**Authors:** Joseph J. Muldoon, Yishan Chuang, Neda Bagheri, Joshua N. Leonard

## Abstract

Macrophage-initiated inflammation is tightly regulated to eliminate threats such as infections while suppressing harmful immune activation. However, individual cells’ signaling responses to pro-inflammatory cues are heterogeneous; subpopulations emerge with high or low activation states, though why this occurs is unknown. To address this question, we used single-cell tracking and dynamical modeling to develop and validate a revised model for macrophage activation by lipopolysaccharide (LPS) that invokes a mechanism we term *quorum licensing*. Bimodal phenotypic partitioning of macrophages is primed during the resting state, depends on cumulative history of cell density, is predicted by extrinsic noise in transcription factor expression, and is independent of canonical LPS-induced intercellular feedback in the tumor necrosis factor (TNF) response. Our analysis shows how this density-dependent coupling produces a nonlinear effect on collective TNF production. We speculate that by linking macrophage density to activation, this mechanism could amplify local responses to threats and prevent false alarms.

---

In responding to external cues, cells are faced with many options, but by sharing information a population of cells can make more effective decisions than can each individual alone^1–3^. These decisions are generally mediated by secreted products. Bacteria use quorum sensing (QS) molecules to coordinate when and whether to form biofilms, and social amoeba secrete cyclic AMP to coordinate their aggregation. In each case, a proxy is used to convey information about the local number of cells available to coordinate the task. In immunology, a canonical example of coordination is the amplification of the response to a perceived threat. During infection, cells such as macrophages provide an immediate line of defense by initiating inflammation to eliminate invading microbes^4^. This process often begins when the bacterial membrane component lipopolysaccharide (LPS), a canonical pro-inflammatory cue, is sensed by Toll-like receptor 4 (TLR4). Prior to TLR4 activation, the transcription factor NF-κB is sequestered in the cytoplasm by Inhibitor of κB (IκB), and upon activation, IκB kinase (IKK) targets IκB for degradation^5,6^, allowing NF-κB to translocate to the nucleus. There, NF-κB induces the transcription of genes such as tumor necrosis factor (TNF), a cytokine that mediates inflammation and pathogen clearance^7–9^. Other pathways downstream of TLR4 increase TNF production by stabilizing the mRNA and promoting translation and cleavage of the proprotein for secretion. Extracellular TNF then signals through TNF Receptor 1 (TNFR1), driving NF-κB activation in positive feedback^10–14^, and this is a general explanation for how macrophages and other cells amplify their response to LPS.

The regulation of TNF has multiple layers. Recently, it was found that when the concentration of LPS exceeds a certain threshold, the induced signaling through NF-κB drives the transcription of its own RelA subunit in a process termed the feedback dominance (FBD) switch, producing *intracellular* positive feedback on NF-κB expression and activity^15^. Other pathways act to constrain the response to LPS and ensure its eventual resolution: cell-intrinsic regulators (*i.e*., those with intracellular origins) include microRNAs and mRNA-binding proteins that decrease *Tnf* stability and translation^16^, as well as IκB^5,17^ and various inhibitors of IKK^6,10,18^ induced by NF-κB in negative feedback; cell-extrinsic regulators (*i.e*., those with extracellular origins) include interleukin 10 (IL-10), in that IL-10 signaling via the IL-10 receptor (IL-10R) antagonizes NF-κB activity and destabilizes *Tnf* stability and translation. In combination, these interlocking positive and negative motifs confer the functional plasticity necessary for immune cells to balance pathogen clearance with harmful side effects such as cytotoxicity and tissue damage^19^.

Given the complexity of the regulation of NF-κB and TNF, computational models have proven valuable for elucidating the dynamical properties of these systems and the roles of individual components. Early models explicated intracellular signaling^5,6,20–22^, and subsequent models included newly appreciated mechanisms such as intercellular feedback^10,12,13,23–26^. Recent studies have incorporated cell heterogeneity, by attributing observed differences in gene expression between cells either to stochastic fluctuations^27–29^ or to variation in initial values^30^, kinetic parameters^15,31–33^, or timing of signaling events^34^. A key consideration for understanding signaling and regulation in macrophages, in particular, is that these cells characteristically exhibit broad phenotypic heterogeneity^35–37^. It has been proposed that this variation could have important functional consequences, *e.g*., to broaden the repertoire of responses to stimuli^38^, to propagate or restrain coordinated actions, or to convert digital single-cell decisions into analog population-level behaviors^39^. While these ideas are intriguing, specific ways in which such heterogeneity may confer functional gain are not well understood.

In this study, we investigate the intriguing observation that when macrophages are treated with LPS, cell subpopulations emerge with high and low activation states. We propose a revised model in which macrophages use a process that we term *quorum licensing* to link the history of their density to the proportion of cells that become highly activated. This investigation provides new insights into how populations of macrophages use density information to regulate their collective activation.

## RESULTS

### TNF expression is heterogeneous and involves intercellular communication

Macrophage phenotypic heterogeneity has been observed in several studies^35–37^, and non-genetic heterogenous activation has been described in the widely used model cell line RAW 264.7^15,35^. Here, we selected the RAW 264.7 model system to investigate how perturbations that modulate the response to LPS affect the heterogeneity with which macrophages become activated, as represented by expression of TNF (**Fig. 1a**). Pre-treatment of cells with IL-10, prior to treatment with LPS, diminished the average intracellular TNF protein expression measured at 3 h post-stimulation (hps), although TNF distributions across IL-10 doses were broad and overlapping (**Fig. 1b, Supplementary Fig. S1a**). Intracellular TNF was not highly correlated with flow cytometric proxies for cell size (**Supplementary Fig. S1b**), suggesting the heterogeneity was not due to cell cycle asynchrony alone.

**Figure 1.**
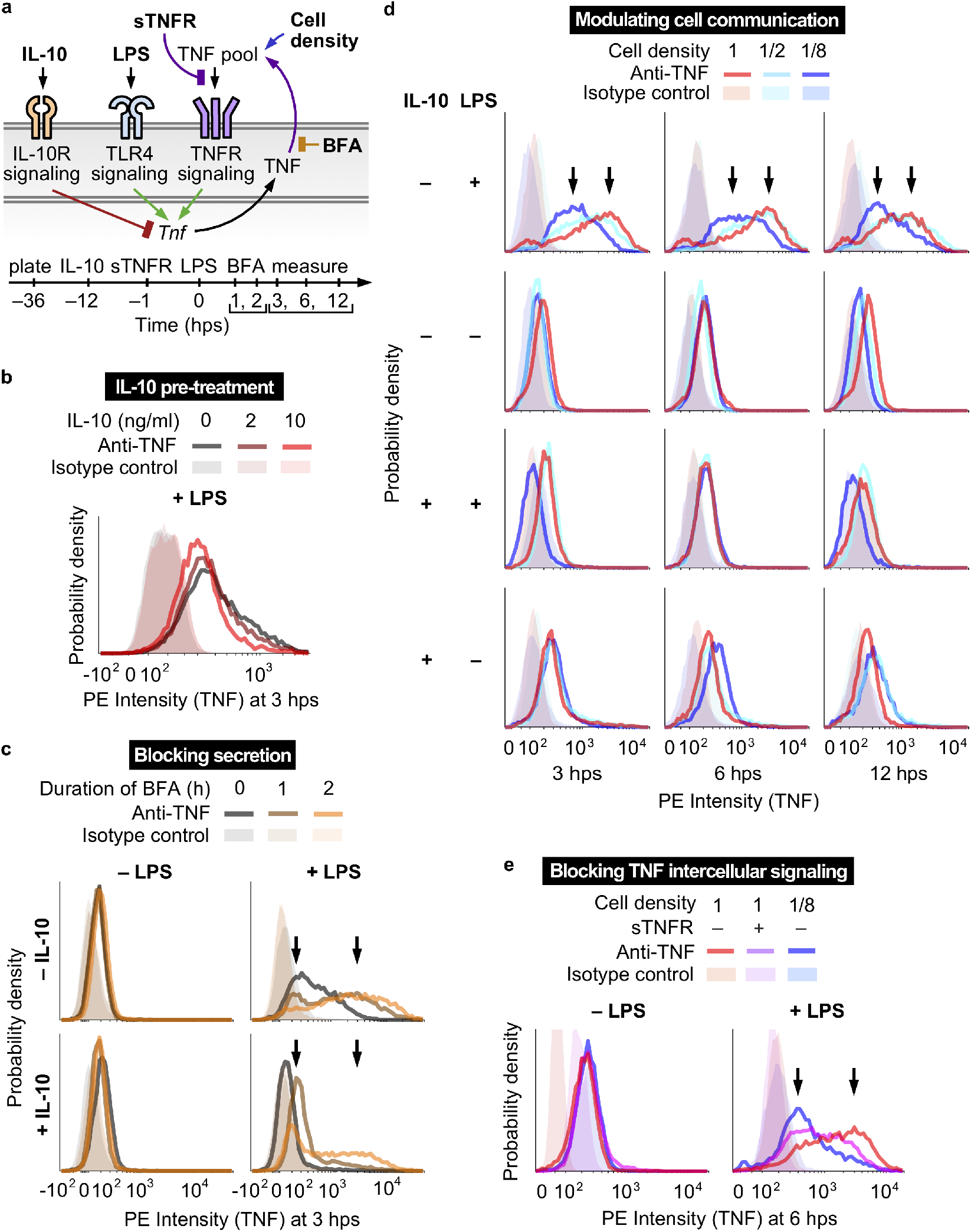
The TNF response to LPS is heterogeneous and requires intercellular communication. (**a**) The diagram summarizes the perturbations and stimuli applied to investigate TNF expression and intercellular communication (hps, hours post-stimulation with LPS). LPS activates TLR4 signaling, which induces TNF expression. IL-10 pre-treatment activates IL-10R signaling, which inhibits LPS-induced TNF expression. Secreted TNF activates TNFR signaling, which induces TNF further through intercellular feedback. BFA prevents cytokine secretion, causing TNF to accumulate intracellularly. Varying the cell density modulates the concentrations of secreted factors such as TNF. sTNFR binds extracellular TNF to prevent TNFR signaling. (**b**) IL-10 pre-treatment diminishes LPS-induced TNF expression. In **b–e**, x-axes are on a logicle scale (linear near zero, and log_10_-scaled farther from zero). (**c**) TNF expression is heterogeneous with high-expressing and low-expressing subpopulations. After pre-treatment with IL-10 (10 ng/ml) and/or treatment with LPS (100 ng/ml), cells were treated with BFA at 1 h or 2 hps. Arrows in **c–e** indicate low and high modes of the TNF distributions. (**d**) The full TNF response to LPS requires intercellular communication. (**e**) Intercellular feedback through secreted TNF is necessary for the full response.

To obtain a more direct readout of TNF production, we applied brefeldin A (BFA) to inhibit anterograde transport from the endoplasmic reticulum to the Golgi apparatus and prevent cytokine secretion^40^. We reasoned that if variation in TNF secretion were a main source of heterogeneity, then BFA would diminish heterogeneity, and if gene regulation were the main source, then BFA would exaggerate it. Without LPS, BFA had no appreciable effect on intracellular TNF, indicating no detectable basal TNF production. However, when added after stimulation with LPS, BFA led to wide-ranging accumulation of on average several-fold more TNF cell^−1^ h^−1^ (**Fig. 1c, Supplementary Fig. S1c**). Unexpectedly, while most cells accumulated more TNF with longer BFA treatment, some cells accumulated little or no TNF over time. When cells were pre-treated with IL-10 prior to LPS, TNF accumulation was less than when treated with LPS only, as expected, yet TNF accumulation was still wide-ranging for a majority of cells and low for a minority. Therefore, blocking secretion unmasked substantial hidden variation in TNF production and showed that the cell population includes both high (wide-ranging) and low responders to LPS, regardless of whether cells are pre-treated with IL-10.

Since LPS-induced TNF paracrine signaling is known to contribute to NF-κB activity^10–13^, we next examined the effect of intercellular communication on TNF production. To modulate communication in a manner that is not biased toward or against specific secreted factors, we varied cell density (full, half, or one-eighth of typical plating conditions) as a general handle for tuning the magnitude of coupling between cells. Without IL-10 and with LPS, a cell density-dependent effect was evident (**Fig. 1d, Supplementary Fig. S1d**). At 3 hps, average TNF expression correlated with density. At 6 hps, half and full density were TNF-high while one-eighth density remained low, and by 12 hps expression had begun to decrease for each case. All of the distributions were heterogeneous, but at high density, intracellular TNF remained skewed toward high expression over time, and at low density, it remained skewed toward low expression. With IL-10 or without LPS, little to no intracellular TNF was detectable. Thus, the full response to LPS requires intercellular communication, with cell density-associated effects on TNF production persisting over time.

To investigate whether secreted TNF sustains its own LPS-induced expression, cells were pre-treated with excess soluble TNF receptor (sTNFR) to titrate extracellular TNF from binding cell surface receptors. As TNF is bound by sTNFR, TNFR signaling is blocked^13^. With LPS, cells at high density with sTNFR pretreatment expressed TNF at an intermediate level on average—less than at high density without sTNFR, and more than at low density without sTNFR—and the distribution remained heterogeneous (**Fig. 1e, Supplementary Fig. S1e**). Therefore, although TNF-mediated intercellular positive feedback is required for full TNF production as expected^10–14^, this mechanism does not account for the wide-ranging TNF expression observed, nor does it explain the high and low activation states observed. Thus, an additional explanation is required.

### The proportion of cells in each activation state depends on cell density

To investigate the phenomena described above, we examined regulation upstream of the TNF protein using a previously validated clonal macrophage cell line with two genomically integrated reporters^15^ (**Supplementary Fig. S2a**). Such a reporter system uniquely enables one to resolve the dynamics and heterogeneity of individual cell signaling responses. The first reporter is a fusion of enhanced green fluorescent protein (EGFP) and RelA (the p65 subunit of NF-κB) driven by the *Rela* promoter. In the resting cell state (pre-LPS), EGFP-RelA is sequestered primarily in the cytoplasm, and upon activation, this protein translocates to the nucleus and induces transcription of NF-κB target genes. Above a sufficient LPS dose, EGFP-RelA also induces the expression of endogenous RelA (and expression of EGFP-RelA) via an intracellular positive feedback loop termed FBD^15^. Thus, EGFP-RelA tracks both the localization and expression of NF-κB. The second reporter is a fusion of mCherry and a destabilizing PEST tag^41^ driven by the *Tnf* promoter. *mCherry* RNA lacks the *Tnf*-specific 3’ UTR and is decoupled from *Tnf*-specific post-transcriptional regulation. These features make the mCherry protein a proxy for transcription from the *Tnf* promoter (rather than downstream TNF protein expression), after accounting for the time delay for mCherry translation and maturation^42^.

We utilized this reporter system to examine RelA expression and localization, and *Tnf* promoter activity, under the perturbations used above (**Fig. 2a, Supplementary Fig. S2b–c**). In all cases without LPS, EGFP-RelA and mCherry expression were low. With LPS, the mCherry distribution shifted from unimodal to bimodal, consistent with the observed expression of endogenous TNF (**Fig. 1c**). At the three high density conditions, most cells were mCherry^high^, and at low density, a greater proportion was mCherry^low^. The outcome at low density was not due to inhibition of EGFP-RelA nuclear translocation (**Supplementary Fig. S2d**). EGFP-RelA mirrored the pattern for mCherry, albeit with more overlap between modes, indicating an association between the reporters’ induction (**Supplementary Fig. S2c**). We also observed that average levels of TNF and mCherry ranked differently across conditions. In particular, “high density +IL-10 +LPS” yielded greater mCherry expression than did “low density +LPS” (**Fig. 2a**), and this ranking was reversed for TNF (**Fig. 1**). The difference could be due to post-transcriptional downregulation of *Tnf* induced by IL-10R signaling^43–46^, which would diminish TNF protein more than *Tnf* transcription, and mCherry is a proxy for the latter.

**Figure 2.**
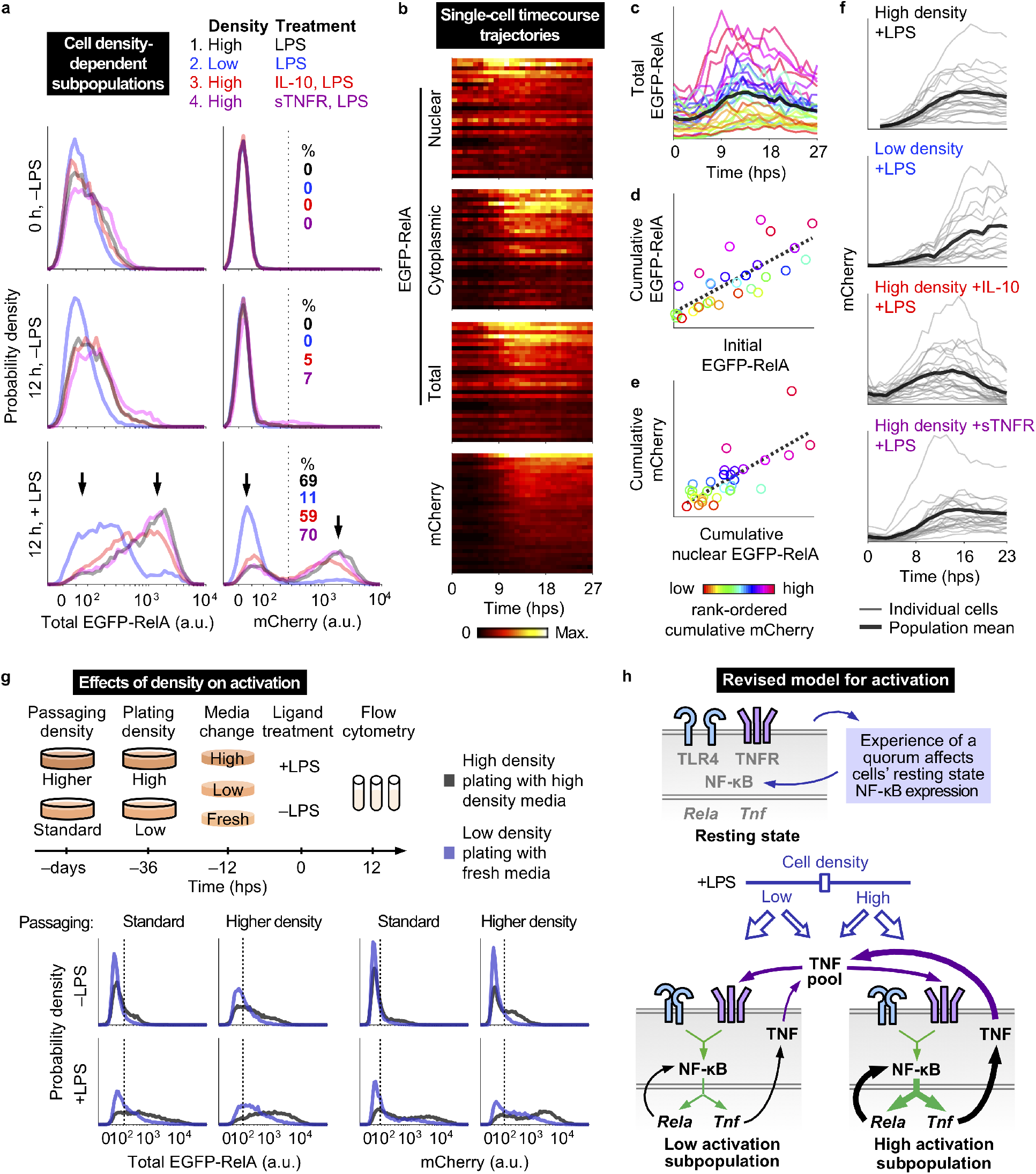
Cell density modulates the heterogeneity of macrophage activation. (**a**) Reporter protein fluorescence was measured by flow cytometry for the indicated cell densities, time points, and ligand treatments (IL-10 and sTNFR treatments as in **Fig. 1**). Percentages of highly activated cells were determined using a threshold (dotted vertical line) at the nadir between the two modes (arrows) of mCherry distributions. (**b**) Reporter trajectories for cells at high density after treatment with sTNFR (1 h pre-LPS) and LPS. Cells (rows) are rank-ordered by cumulative mCherry expression and color-coded by fluorescence magnitude within heat maps. (**c**) Single-cell trajectories of total EGFP-RelA expression are color-coded red-to-magenta, for low-to-high rank-ordered cumulative mCherry expression. The population mean is in bold. (**d–e**) Relationship between initial and cumulative total EGFP-RelA (R^2^ = 0.61, one-tailed permutation test *p* = 2×10^−7^) and between cumulative nuclear EGFP-RelA and cumulative mCherry (R^2^ = 0.59, one-tailed permutation test *p* = 3×10^−7^). Dotted lines are linear fits; axes are linearly scaled. (**f**) Single-cell mCherry trajectories, with the population mean in bold. Values are in a.u. specific to each subpanel. (**g**) Effect of culture density-associated conditions on macrophage activation and heterogeneity. Fluorescence units are comparable within each reporter protein and passaging density. Vertical lines are to guide the eye to low and high activation states. (**h**) Revised conceptual model for macrophage activation with differently activated cell density-dependent subpopulations.

Altogether, macrophage activation was bimodal, with subpopulation proportions that varied with cell density. This observation held under perturbations to TNF-regulating pathways such as TNFR and IL-10R signaling. The dependence of the decision to become highly activated on cell density has some resemblance to bacterial quorum sensing (QS), in that the phenotypes of individual cells are determined by information shared by the population. However, an important distinction is that while in QS essentially all cells become activated if a threshold concentration of QS molecule is surpassed^47^, here the *proportion* of highly activated cells increases with density. At the population level, this relationship produces an analog response rather than a digital one. To distinguish these two phenomena, we refer to the macrophage behavior as *quorum licensing*.

### Single-cell analysis of RelA and *Tnf* dynamics

We next investigated whether heterogeneous regulation of activation was due to variation in the magnitude and/or timing of the response to LPS. Variation in the magnitude of the response would indicate a role for intrinsic or extrinsic noise, due to stochastic fluctuations or to deterministic outcomes of variation in initial (pre-LPS) conditions, respectively. Variation in the timing of the response could indicate a domino effect where early high-expressing cells activate other cells. To track the dynamics of EGFP-RelA localization and expression and mCherry expression, cells were stimulated and monitored over a one-day timecourse using confocal laser-scanning microscopy. Quantification of individual trajectories using image analysis software^48^ showed that *Tnf* promoter activation varied primarily in magnitude rather than timing (**Fig. 2b**). For EGFP-RelA (at high density with sTNFR and LPS), the mean expression increased and peaked at 14±4 hps (± standard deviation among cells), after the peak nuclear intensity (10±6 hps) and near the time of peak cytoplasmic intensity (15±4 hps). The magnitude of EGFP-RelA peak intensity was greater in the cytoplasm than the nucleus (1.4±0.4 fold), and depletion of the nuclear portion mid-timecourse coincided with cytoplasmic accumulation. Expression of mCherry was initially low, and it increased and peaked at 18±5 hps. A small subpopulation of cells expressed very little mCherry or EGFP-RelA (consistent with the earlier flow cytometry observations for TNF and the reporters), and reporter expression appeared unrelated to the number of cells in physical contact with one another (**Supplementary Fig. S2e**). Intriguingly, although high induction of both reporters co-occurred in the same cells, the bimodality in TNF expression (measured at 3 hps in **Fig. 1c**) was evident *before* the observed increase in EGFP-RelA fluorescence (**Fig. 2b**), even considering the ~1 h chromophore maturation time. This sequence of events was also observed for cells cultured at low density (**Supplementary Fig. S2f**). Since target gene expression would be expected to increase *after* an increase in the expression of a transcriptional regulator for that gene, the observed sequence indicates that the RelA FBD switch cannot be causal for the early TNF burst observed. Instead, these events are conditionally independent, *i.e*., regulated by an upstream process, and not by each other.

The microscopy analysis revealed additional features of the single-cell responses. For nuclear-localized EGFP-RelA, while the average profile underwent an overall increase and subsequent decrease, some cells exhibited multiple peaks with varied amplitudes (**Supplementary Fig. S2g–h**) in agreement with a recent study^33^. There was also variation in an apparent nuclear reservoir of transcription factor (**Fig. S2i–j**), in agreement with another study^32^. For total EGFP-RelA, the pre-LPS expression varied and was correlated with the post-LPS cumulative expression (**Fig. 2c–d, Supplementary Fig. S2k**). The predictive power of this initial condition indicates a role for extrinsic noise (*i.e*., variation initially present in the system) in determining the post-FBD amount of transcription factor. Cumulative nuclear EGFP-RelA and cumulative mCherry were also correlated, as expected for the pairing of a transcription factor and a target promoter (**Fig. 2e**). These outcomes indicate that extrinsic noise in RelA expression propagates to activity at the *Tnf* promoter. To further examine promoter activity, we quantified mCherry trajectories under each perturbation (**Fig. 2f**). At high density with LPS, mCherry increased for 16±2 hps, indicating continued transcription after measurements of intracellular TNF protein showed a decrease (**Fig. 1d**). This protein decrease is consistent with known post-transcriptional mechanisms that eventually downregulate *Tnf*^16^, and such simultaneously opposing trends of transcriptional and post-transcriptional regulation represent a type of control that has been likened to operating both the throttle and brake pedals of a vehicle^49^. Cells at low density took more time to reach peak mCherry intensity post-LPS compared to cells growing at high density. Cells pretreated with IL-10 increased in mCherry for 12±3 hps and then decreased toward basal levels. Cells treated with sTNFR responded to LPS similarly to cells without this antagonist. Together, the measurements from flow cytometry and microscopy show how TNF is differently regulated under each perturbation (summarized in **Supplementary Fig. S2i**).

An additional phenomenon related to cell density is that EGFP-RelA levels differed between low and high density conditions, both with and without LPS stimulation (**Fig. 2a**). This difference suggests that secreted factors might affect cells’ resting states in a way that predicts the response to LPS. To more carefully investigate how exposure to high density culture-associated conditions impact the patterns described above, we exposed cells to combinations of: standard or higher density passaging (for at least three days), low or high density plating (at 36 h pre-LPS), and exposure to different conditioned media (fresh, low, or high; at 12 h pre-LPS). In general, exposure to more high density-associated conditions increased the basal and inducible reporter expression (**Fig. 2g, Supplementary Table S1**). For cells that were passaged at higher density and later provided conditioned media from high density cells, the outcomes were similar regardless of cell density at plating (which determines the number of cells present), indicating that sustained exposure to high density-associated secreted factors was sufficient to prime and enable full LPS-induced activation regardless of cell density (**Supplementary Fig. S2m**). For the highest-density combination of conditions, reporter bimodality was prominent and right-shifted, even prior to LPS stimulation (**Fig. 2g**). The effect of passaging (which took place over several days prior to plating, which was itself 36 h pre-LPS) on the distributions shows it is heritable across cell generations. These observations motivate a revised model for macrophage activation in which factors secreted during the resting state modulate the propensity for cells to become highly activated (**Fig. 2h**). The number of cells and the density-dependent proportion that become highly activated have a multiplicative effect on TNF production.

### A new computational model for heterogeneous macrophage activation

To integrate our observations with prior knowledge on macrophage activation, we developed a computational model for the intracellular and intercellular signaling network. We reasoned that a dynamical model could elucidate how TNF is regulated and the role of cell heterogeneity. Such a framework could enable us to investigate whether heterogeneity confers advantageous properties to a population, and how different regulatory influences on TNF are coordinated over time. Given the complexity of this system, key aspects of model development^50^ were to include the most essential components, formulate equations to concisely portray biochemical processes, identify salient features of the data, calibrate parameters, and evaluate the extent to which model simulations could explain the experimental observations.

To start, we examined prior experimental and computational studies on NF-κB signaling and TNF regulation^12–15,21,26,32,33,43–45,51–56^ (**Supplementary Information**) and synthesized this information to produce a system of ordinary differential equations (ODEs) representing a cell that can inducibly express, secrete, and sense TNF. Model reduction principles were applied to decrease complexity^57^. For instance, we removed non-rate limiting steps and non-influential variables. Then, through several iterations, various network topologies and corresponding model formulations (sets of equations) were proposed, evaluated, and refined. Since extrinsic noise featured prominently in the data, we hypothesized that it might be possible to first calibrate a homogeneous model (*i.e*., a model of one cell) based on mean-average flow cytometry and microscopy data, and then incorporate heterogeneity at a later point. Calibration was conducted using parameter sweeps, multi-objective optimization, and an evolutionary algorithm to score, cull, repopulate, and mutate parameter sets over many generations (**Supplementary Information**).

The nonlinearity of the system combined with high dimensionality of the search space presented a challenge for calibration, so we took a strategy of splitting the parameters into two groups to be fitted in separate rounds. The first round used a cell-intrinsic model of NF-κB activation, which covered TLR4 signaling; NF-κB activation, nucleocytoplasmic translocation, and inactivation; IκB expression; and the FBD switch (**Supplementary Fig. S3a–b**). A fit to data for sTNFR and LPS (to exclude the influence of TNFR) resulted in a family of parameter subsets yielding similar outcomes. The second round of parameter estimation used the full model, which incorporates *Tnf* transcriptional, post-transcriptional, and translational regulation; mCherry expression; effects of IL-10 pre-treatment on NF-κB and TNF expression (at multiple stages of regulation); TNF secretion and its inhibition by BFA; the effect of population growth on extracellular TNF (using time-dependent logistic growth, **Supplementary Fig. S3c**, noting that neither LPS nor IL-10 affected cell viability); and TNFR signaling. Experimental perturbations (**Fig. 1–2**) were simulated by modifying the equations to capture each perturbation’s mode of action (**Supplementary Fig. S3d**). A simultaneous fit to all of the data, in which some parameters were constrained to the subsets from the first round and the others underwent a free search, yielded the homogeneous model (depicted in **Fig. 3a, Supplementary Tables S2–S5**).

**Figure 3.**
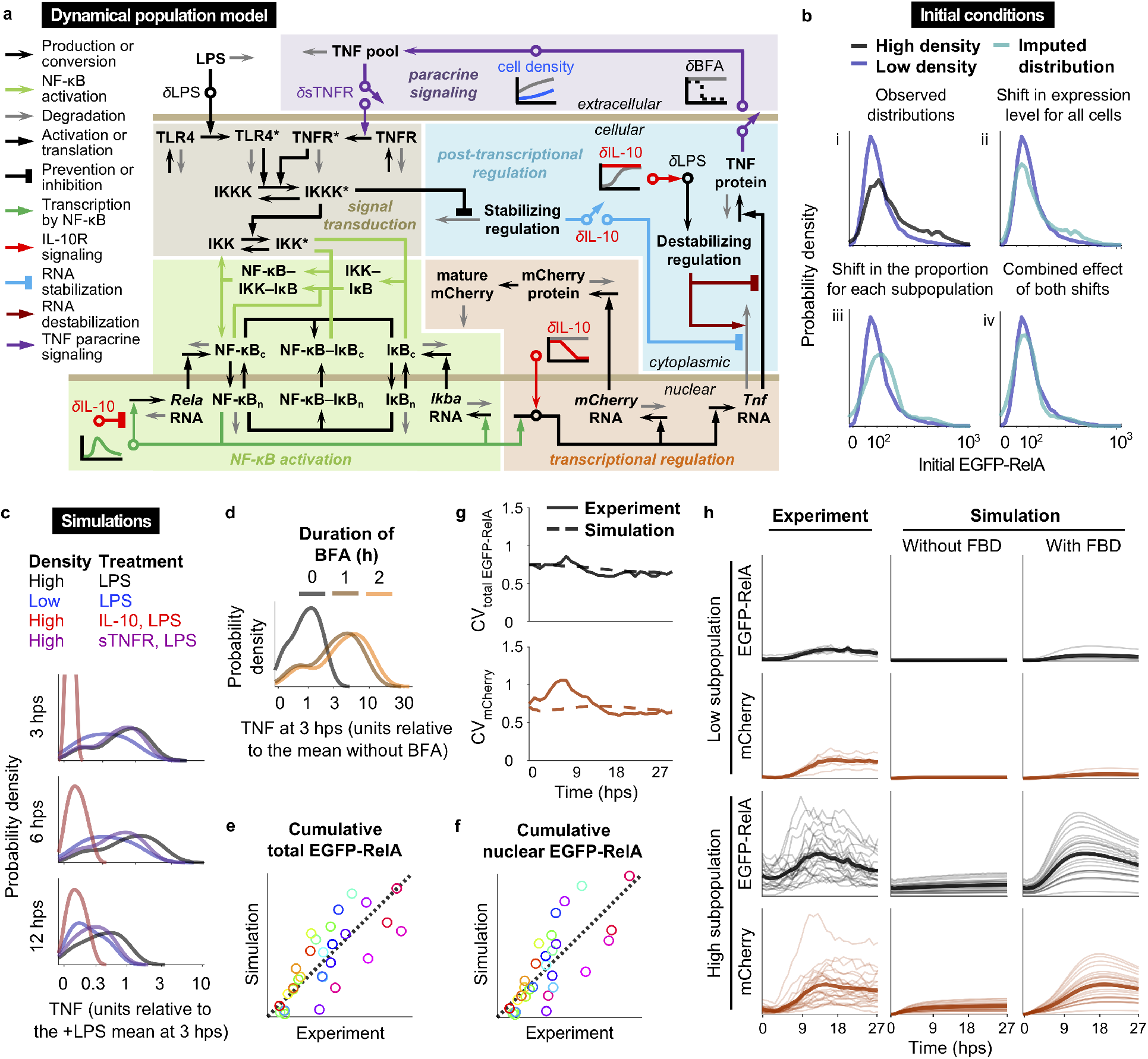
Dynamical population model for the signaling and regulatory network. (**a**) The diagram summarizes the variables, reactions, and mechanisms in the model of macrophage activation with quorum licensing. Symbols: bold non-italicized text for variables; horizontal bars for compartment boundaries; n for nuclear and c for cytoplasmic; asterisks for activated receptors or kinases; δ and circles for perturbation-specific effects; diagonal arrows for ON/OFF effects; graphs for time-dependent processes. (**b**) Generation of comparable distributions of initial values. The observed distributions in (i) are in non-comparable units; to initiate simulations with NF-κB distributions that vary between high and low cell density and that match experimental observations for EGFP-RelA, we impute from an observed high density distribution (black) a low density distribution (teal) that matches the observed low density distribution (blue). The transform shifts the distribution (ii) and adjusts the proportion of cells in high vs. low states using a Gaussian mixture model of two populations fit to the high density distribution (iii), such that the combined transform (iv) generates an imputed distribution matching the observed low-density distribution (blue) with units relatable to high density (black). (**c–d**) Simulated TNF distributions from the calibrated model match experimental trends for cell density and ligand conditions (**Fig. 1d–e**) and BFA conditions (**Fig. 1c**). All cells have the topology in **a** and are heterogeneous in the initial value and transcription of *Rela* RNA and the initial value of cytoplasmic NF-κB–IκB, derived from **b**. To approximate logicle scaling characteristic of flow cytometry data and to prevent stretched scaling approaching log zero, simulated outcomes were plotted after adding a constant value uniformly and log-transforming the values. (**e–h**) Comparison of simulated and experimental outcomes for cells at high density with sTNFR and LPS. (**e–f**) Cumulative total EGFP-RelA (R^2^ = 0.61, one-tailed permutation test *p* = 3×10^−7^) and cumulative nuclear EGFP-RelA (R^2^ = 0.60, one-tailed permutation test *p* = 4×10^−7^). Dotted lines are linear fits; axes are linearly scaled. (**g**) Coefficient of variation (CV) in mCherry and total EGFP-RelA expression over time. (**h**) Trajectories were grouped *post hoc* by high or low activation. Mean averages are in bold. Simulations are shown with and without FBD.

We next adapted this model to represent a population of cells that are coupled by paracrine feedback, and in which each cell has the same topology but varies in extrinsic noise in NF-κB. To this end, initial values for the NF-κB variable were assigned based on EGFP-RelA measurements from confocal microscopy (**Fig. 2d**). Additionally, we observed that EGFP-RelA distributions differed between high and low density cultures by a shift in values and a shift between the two activation modes (**Fig. 3b**), and applying a transformation *in silico* to the initial values at high density could produce a distribution in comparable units for initial values at low density. In summary, the model was trained on *homogeneous post*-LPS data and initialized with *heterogeneous pre*-LPS data.

Remarkably, the use of varied EGFP-RelA levels prior to LPS stimulation, and the transformation of this distribution between high and low density conditions, enabled the resulting simulations to accurately capture heterogeneity in the data post-LPS, including intracellular TNF expression across perturbations and over time (**Fig. 3c**, compare to **Fig. 1d–e**) and accumulation with BFA (**Fig. 3d**, compare to **Fig. 1c**). These simulations also accounted for the majority of the variation in cumulative transcription factor expression and localization (**Fig. 3e–f**). Furthermore, they closely tracked the distributions and trajectories of reporter expression (**Fig. 3g–h**), and supported that cells in both the high and low activation subpopulations underwent FBD (**Fig. 3h**). These findings support our strategy of first training the model on population average data and then introducing cell heterogeneity in a way that incorporates the effects of cell density to capture how cells’ pre-LPS state predicts the response to LPS.

### Model validation on a distinct test dataset

We next sought to validate the model by testing whether it could predict responses to conditions not included in model development and parameter estimation. To this end, we simulated populations of cells growing at different densities and treated with varied doses of LPS. The FBD switch was included for conditions corresponding to LPS doses at or above 1 ng/ml (our estimate for a threshold at which this mechanism is active, based on the original study^15^). To obtain an analogous test dataset, reporter expression was measured by flow cytometry at 12 hps (**Fig. 4a, Supplementary Fig. S4a**). We observed that the model predictions were broadly in agreement with the experimental data: distributions for EGFP-RelA and mCherry had more rightward shifts with higher LPS doses, and two subpopulations were present with density-dependent proportions across doses. At lower doses and densities, the characteristic high-activation subpopulation had diminished reporter expression, resulting in more overlap between subpopulations within each distribution.

**Figure 4.**
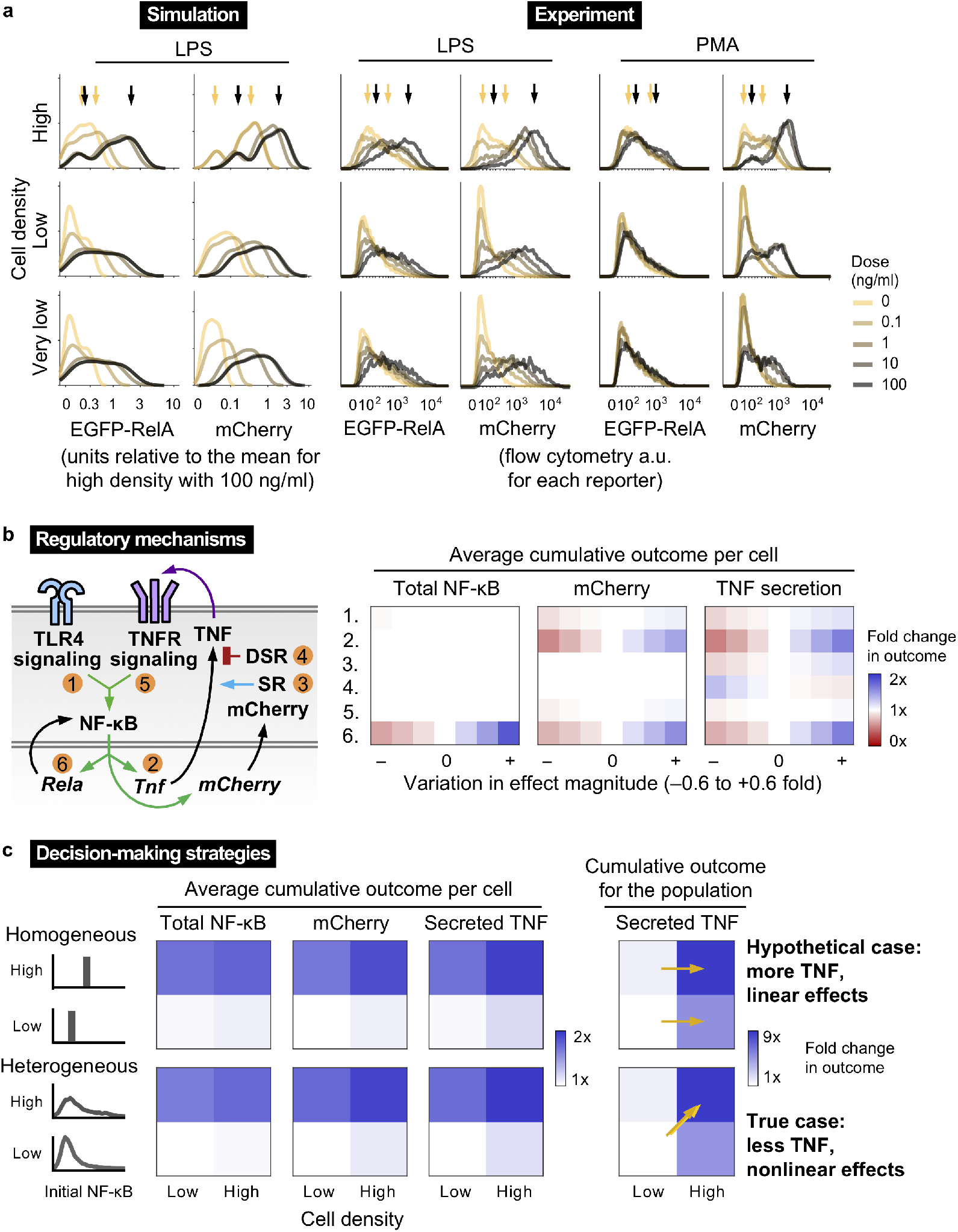
Phenotypic and functional consequences for mechanisms of TNF regulation. (**a**) Predicted and measured distributions of reporter expression across cell densities and doses of LPS or PMA at 12 hps. Very low density is 1/8th of low density. Arrows indicate the modes of the distributions for high density without stimulus or with 100 ng/ml of LPS or PMA. X-axes are on a logicle scale for flow cytometry and transformed for simulations as previously (**Fig. 3**). For simulations: low and very low density used the same initial values; mCherry distributions without stimulus were obtained without FBD and without paracrine feedback; and FBD was applied for LPS doses at and above 1 ng/ml. (**b**) Simulated outcomes after individually varying the effect magnitudes of mechanisms that regulate TNF, with perturbations numbered as depicted in the diagram. Outcomes were assessed using the 12 hps time-integrated amounts of total EGFP-RelA, mCherry, and secreted TNF (*i.e*., cumulative secreted flux) per cell at high density. Conditions corresponding to the origin on the x-axis indicate base case effect magnitudes (*1x* on the color scale for outcomes). Abbreviations: stabilizing regulation (SR), destabilizing regulation (DSR). (**c**) Comparison of activation for homogeneous (hypothetical) or heterogeneous (observed) initial distributions of transcription factor. For each readout, a value of *1x* was set for the outcome given a heterogeneous population at low density with low initial values. Homogeneous initial values were set at the mean averages of corresponding heterogeneous distributions. In the right panel, the effect of the number of cells is incorporated into the total amount of secreted TNF. Arrows indicate relevant comparisons, with key trends noted in bold text.

To test if any of the above observations were specific to TLR4 signaling, we evaluated the same conditions, but exchanged LPS for phorbol 12-myristate 13-acetate (PMA), a membrane-permeable small molecule that activates protein kinase C (PKC), which then activates IKK and by extension NF-κB. As was observed with LPS stimulation, PMA-induced expression of both reporters was bimodal with cell density dependence, and mCherry expression was ligand dose-dependent (**Fig. 4a**). Therefore, quorum licensing-associated activation of the *Tnf* promoter, via NF-κB, does not require that NF-κB be activated via TLR4. Interestingly, unlike trends observed with LPS, RelA expression following PMA addition differed little from the case without PMA, which indicates that FBD is induced by NF-κB activation via TLR4 but not via PKC.

### Model analysis elucidates phenotypic and functional roles for TNF regulatory mechanisms

We next employed the validated model to examine the roles of different TNF regulatory mechanisms. As a first step, we evaluated the robustness of simulations to global parameter variation, by sampling values for the fitted parameters from sets of distributions with different coefficients of variation (CV) centered on the fitted values. Variation was tolerated to some extent (**Supplementary Fig. S4b**), which is consistent with the general phenomenon of *sloppy* parameter sensitivities in systems biology models^58^, generally indicating that a range of values for any one parameter or combination of parameters can yield similar fits to available data. However, once parameter values were varied beyond a CV of ~0.1–0.2, outcomes began to noticeably diverge, suggesting the system might be sensitive to certain mechanisms. We then used the model as a testbed to explore how key behaviors may depend upon the quantitative characteristics of individual mechanisms. To that end, the effect magnitudes of individual mechanisms were varied in the model, and the consequences were evaluated by quantifying relevant simulated outcomes: the mean average cumulative amounts of NF-κB and mCherry (phenotypic consequences, as they report on intracellular regulation) and of TNF secretion (a functional consequence, as it intercellularly coordinates the response to LPS). Of the mechanisms evaluated, FBD was the only one to which the NF-κB readout was sensitive, and all three readouts were strongly affected by FBD (**Fig. 4b, Supplementary Fig. S4c**). Expression of mCherry and TNF were both sensitive to transcriptional induction by NF-κB, as expected. TNF expression had the highest sensitivity to stabilizing regulation, destabilizing regulation, and TLR4 signal transduction. Interestingly, TNFR signal transduction arising from intercellular feedback affected mCherry and TNF expression only modestly, suggesting that even if cells had greater signal transduction efficiency, there would not be a large functional gain. Overall, the analysis indicated that the most consequential mechanisms in the system behaviors studied were FBD and transcriptional induction by NF-κB.

Lastly, we used the model to investigate whether the observed cell heterogeneity could produce outcomes that differ from a hypothetical homogeneous population. Specifically, we examined scenarios of high and low cell density in combination with high and low resting state levels of NF-κB. Homogeneous cells were defined with the mean-average value (*i.e*., without extrinsic noise) from the corresponding heterogeneous distribution, and LPS-induced outcomes were assessed using the same readouts as in **Fig. 4b**, along with the total TNF secreted by the population. Unexpectedly, across scenarios, the homogeneous population was slightly more activated and secreted more TNF than did the heterogeneous population (**Fig. 4c** left panel, **Supplementary Fig. S4d**). Within both populations, for individual cells, the initial conditions had a larger effect on the readouts than did the number of cells, as depicted by a greater distinction between rows than columns in the heatmaps. For the total TNF secretion, both of these factors were important, and the coupling of initial NF-κB to cell density produced a greater fold change in TNF than would be the case if these factors were independent (**Fig. 4c** right panel, diagonal arrow vs. horizontal arrows). This coupling was also more consequential than the distinction between population heterogeneity and homogeneity. Therefore, we conclude that it is not heterogeneity per se that is important for decision-making, but instead that the heterogeneity is regulated in a density-dependent manner leading to bimodal activation.

## DISCUSSION

This study explores the observation that within a clonal population of genetically identical macrophages, a potent activating stimulus (LPS) drives high expression of TNF in only a subset of cells. The observation of TNF heterogeneity is consistent with a recent study that showed cellular responses vary widely in response to such cues^59^. In examining the regulation that underlies this variation, we observed heterogeneity in transcription factor expression and localization, *Tnf* promoter activation, and TNF expression. Furthermore, the measured distributions showed states for high (wide-ranging) and low (approximately off) activation. This bimodality was tunable by modulating conditions related to cell density, indicating a role for intercellular communication potentially via secreted proteins, metabolites, or extracellular vesicles^60^. Using single-cell tracking and dynamical modeling, we proposed, developed, and validated a revised model for activation by *quorum licensing*, in which a population’s experience of density over time impacts the extent to which cells become poised for activation.

Although bacterial QS provides a useful conceptual reference point, quorum licensing more closely resembles other types of phenotypic bimodality in genetically uniform populations, which have been found in native and synthetic contexts, across species, and can stem from various mechanisms. Some of these mechanisms do not require intercellular communication. For example, continuous variation in protein expression in a population has been shown to produce digital outcomes for growth factor-induced kinase activation in individual cells due to threshold effects^61^, and synthetic positive feedback on a promoter with multiple binding sites has been shown to support stochastic activation through transcriptional bursting^62^. Other instances of phenotypic bimodality do involve intercellular communication. For example, T cell secretion of the cytokine IL-2 feeds back through IL-2R signaling to promote further IL-2 expression. Since the sensing step involves capturing extracellular IL-2, the feedback is confined to local regions and only some cells sense sufficient to IL-2 to proliferate^63^. Competitive cytokine uptake is involved in TNF paracrine signaling as well^64^. In the case of a synthetic genetic circuit implemented in yeast, with topological similarity to TNF intercellular feedback and that exhibited strong positive feedback and rapid degradation of the secreted cue, the proportions of cells in upper and lower modes of activation were found to depend on cell density^65^. Mathematical modeling showed that if there is increased reliance on autocrine signaling at low density compared to high density, and if at low density few cells can activate through autocrine signaling alone, then at high density more cells should activate as the balance shifts toward paracrine signaling where a cue is shared. Quorum licensing could belong to a family of bimodal activation behaviors, which may comprise a general phenomenon—many cytokines are regulated by pathways that overlap with those that regulate expression of TNF^45^, and single-cell RNA-seq analysis has revealed bimodality in hundreds of immune response gene transcripts^66,67^. In addition, macrophages were recently found to use a quorum-based mechanism to *resolve* inflammatory processes via production of nitric oxide^68^. Investigating how various modes of heterogeneity and coordination interact represents an exciting avenue for future investigation.

Our study suggests nuanced interpretations for several recently identified phenomena. First, it was shown that NF-κB can induce RelA in intracellular positive feedback termed the FBD switch^15^. We found that this effect persists over the long-term and coincides with sustained *Tnf* promoter activity. However, these dynamics differ from those of intracellular TNF protein, which undergoes a burst (in which bimodality in its production is evident) before FBD takes effect, and then begins to decline before mCherry reaches peak expression. Second, phenotypic variation among genetically identical cells is often attributed to stochasticity (intrinsic noise). However, in this system, the predictive power of the pre-LPS state indicates a substantial role for extrinsic noise. If additional species were measured to more fully characterize the pre-LPS state, we anticipate that the multivariate initial conditions would provide further predictive power as to how individual cells are poised to respond to LPS. Third, the role of TNF in intercellular communication has been widely studied^10–14^. However, we found that as an intercellular signal, TNF had a modest influence on FBD and *Tnf* promoter activity. Intercellular TNF signaling had a larger effect on TNF protein expression, consistent with known mechanisms through which TNFR signaling enhances TNF expression through post-transcriptional regulation^54^. Since chronic TNF production is implicated in various diseases, therapeutic strategies have focused on blocking TNF from binding surface receptors by administering anti-TNF antibodies or soluble TNFR^69^, or administering antagonists of TLR4 or associated proteins^70^. While we did not identify specific molecules that mediate the observed quorum licensing, if these factors can be identified in future work, they may represent additional targets for immunomodulation.

Looking beyond the scope of this investigation, it is interesting to speculate whether quorum licensing may be adaptive for immune function. Such a mechanism could provide a way for a cell population to *preemptively* coordinate a response to microbial incursion prior to LPS-induced intercellular signaling. As an example, we consider a scenario in which a wound is experienced, resulting in microbial incursion. Since tissue damage can immediately trigger local sterile inflammation (via damage-associated molecular patterns, DAMPs)^71^, macrophages among other cells could locally accumulate and become primed for high activation through quorum licensing. Then, should replication of invading microbes produce more pathogen-associated molecular patterns (PAMPs) such as LPS, the local inflammatory response would escalate. Another consequence would be to limit potent inflammation to local environments. In the scenario posed above—a wound experiencing an infection—some DAMPs may travel to sites that are remote from the wound, even if the microbes are limited to the wound site. Quorum licensing could act to limit the most potent macrophage-mediated responses to sites characterized by *both* the presence of DAMPs and the local accumulation of macrophages, nonlinearly driving local cytokine production. If macrophage recruitment could be coupled to activation in this way, then quorum licensing would enhance the specificity of this potent but potentially harmful facet of innate immune function. Conversely however, for conditions such as sarcoidosis, fibrosis, and atherosclerosis^72,73^ characterized by abnormal accumulation of macrophages and related cells, quorum licensing would be maladaptive by supporting chronic inflammation. These ideas each comprise compelling avenues for future investigation, building upon the insights gained here into mechanisms linking heterogenous macrophage activation to intercellular communication.

## METHODS

### Cell culture

RAW 264.7 cells were cultured in complete DMEM medium (containing DMEM, 10% heat-inactivated FBS, and 4 mM L-glutamine) in tissue culture-treated dishes (Corning). A typical plating prior to a flow cytometry experiment used 1.35 ml of cell culture per well of a 6-well plate (3.3×10^5^ cells/ml for high density, 4.1×10^4^ cells/ml for low density). To obtain sufficient cells at very low density (**Fig. 4**), 8 ml of cell culture was used in a 10 cm dish (5.2×10^3^ cells/ml). Cells were treated as indicated with IL-10 at 12 h pre-LPS (10 mg/ml, except as otherwise indicated, **Fig. 1b**), recombinant mouse sTNFR (8.3 μg/ml, R&D Systems) at 1 h pre-LPS, *E. coli* 055:B5 LPS (Sigma-Aldrich) or phorbol myristate acetate (Cayman Chemical) at the 0 h time point (36 h after plating), or brefeldin A (Sigma-Aldrich, 2 μg/ml) at 1 or 2 h post-LPS. Media was not changed during the span of ligand treatments. For passaging, cells were incubated briefly in PBS-EDTA (5 mM EDTA in PBS pH 7.4), suspended by gentle scraping, pelleted by centrifugation (125×g, 5 min), and resuspended in medium and plated. For the higher density passaging (**Fig. 2**), cells were grown to cover a large majority of surface area of the dish prior to passaging, and this condition was maintained for several days leading up to the experiment. RAW 264.7 cells were a gift from David Segal (NIH), and reporter cells were a gift from Iain Fraser (NIH)^15^.

### Fixing samples

Media was aspirated, and cells were incubated in PBS-EDTA, detached from the plate, and pelleted. Supernatant was decanted, paraformaldehyde (4% PFA in PBS, 30 μl) was added, and samples were incubated at 4°C for 20 min. PBS-EDTA was added, cells were pelleted, and supernatant was decanted; this step was repeated. Several drops of flow buffer (5 mM EDTA and 0.1% BSA in PBS) were added, and samples were stored at 4°C until staining or flow cytometry.

### Immunostaining

RAW cells were fixed and then suspended in permeabilization wash buffer (0.5% saponin and 0.2% BSA in PBS). Surface and intracellular non-specific binding was blocked using mouse serum. PE-conjugated rat anti-mouse TNF antibody (BD Bioscience #554419) was added to mouse serum-containing cell samples. Cells were incubated at 4°C for 1 h, washed two times with flow buffer, and collected for flow analysis. Isotype control samples used a PE-conjugated rat isotype control antibody (BD Bioscience #554685).

### Flow cytometry

Samples were run on a BD LSRII or BD LSR Fortessa flow cytometer, using the FITC parameter for EGFP-RelA and the PE-Texas Red parameter for mCherry. Data were analyzed using FlowJo software. For experiments with reporter cells, compensation controls were made by transfecting HEK293FT cells with plasmids for constitutive EGFP or mCherry expression. Samples were gated on live (FSC-A vs. SSC-A) and single-cell (FSC-A vs. FSC-H) bases, and fluorescence values were exported from FlowJo and imported into Matlab for further analysis.

### Confocal microscopy

Experiments were conducted using a Zeiss inverted Axio Observer Z1 confocal microscope within a custom environmental control chamber for CO_2_, humidity, and temperature control. Fluorescence intensity was quantified using ImageJ software. At each time point, the mean background pixel intensity (based on multiple sampled regions) was subtracted from the mean intensity of each cell, and this value was multiplied by the area enclosed by the plasma membrane in the 2-D image slice to yield the total intensity in arbitrary units (a.u.). A similar method was applied to the area enclosed by the nucleus to yield the nuclear intensity for EGFP-RelA. At high density, 30 cells were quantified. Cells were excluded from quantification if they divided or exited the field of view during the timecourse. At low density, 20 cells were quantified; due to the low number of cells that could be tracked within the field of view, some traces start or end within the timecourse.

### Visualization

Flow cytometry data were plotted as probability distributions with a logicle x-axis, a standard method that uses linear scaling near the zero mark and log10-scaling farther from zero^74^. Confocal trajectories were plotted with a linear y-axis. For simulations corresponding to flow cytometry, to approximate logicle scaling and facilitate comparisons with experimental data, a small arbitrary constant value was added uniformly to the simulated values, and probability distributions were plotted with a log_10_ x-axis.

### Statistical analysis

*P*-values for **Fig. 2d–e** and **Fig. 3e–f** were determined using one-tailed permutation tests (*n* = 30 cells).

### Code availability

Model formulation, development, and calibration is discussed in **Supplementary Information**. Upon publication, model files will be made available online at github.com/bagherilab.

### Data availability

Data are available in supplementary information files and from the corresponding authors upon request.

## Acknowledgements

The project described was supported by a Cornew Innovation Award from the Chemistry of Life Processes Institute at Northwestern University (J.N.L., N.B.). This work was supported by the Northwestern University Physical Sciences-Oncology Center (http://www.psoc.northwestern.edu) through National Institutes of Health award U54 CA143869-05 (J.N.L., N.B.); National Institutes of Health Award 1R21AI131179-01A1; startup funds from Northwestern University (J.N.L., N.B.); and the Northwestern University – Flow Cytometry Core Facility supported by Cancer Center Support Grant (NCI CA060553). We thank Daniel Wells, William Kath, and Aushra Abouzeid for guidance on the model; Keith MacRenaris, Michelle Hung, Kelly Schwarz, Taylor Dolberg, Patrick Donahue, and Devin Stranford for guidance on experiments; Sean Allen for reagents; and Sarah Stainbrook, Gokay Yamankurt, Patrick Donahue, and members of the Bagheri lab and Leonard lab for useful discussions.

## Author contributions

Conceptualization, J.J.M., Y.C., N.B., J.N.L.; Methodology, J.J.M., Y.C., N.B., J.N.L.; Software, J.J.M.; Validation, J.J.M., Y.C.; Formal analysis, J.J.M., Y.C.; Investigation, J.J.M., Y.C.; Resources, J.N.L.; Data curation, J.J.M.; Writing – original draft, J.J.M.; Writing – review & editing, J.J.M., Y.C., N.B., J.N.L; Visualization, J.J.M.; Supervision, N.B., J.N.L.; Project administration, N.B., J.N.L.; Funding acquisition, J.J.M., N.B., J.N.L.

## Additional information

**Supplementary information** accompanies this paper.

## REFERENCES

1 Perkins, T. J. & Swain, P. S. Strategies for cellular decision-making. Mol Syst Biol 5 (2009).

2 Helikar, T., Kochi, N., Konvalina, J. & Rogers, J. A. in Systems biology for signaling networks Ch. 12, 295–336 (Springer-Verlag, 2010).

3 Shin, Y.-J. & Mahrou, B. in Conf Proc IEEE Eng Med Biol Soc 334–337 (IEEE, Chicago, IL, 2014).

4 Liu, G. & Yang, H. Modulation of macrophage activation and programming in immunity. J Cell Physio 228, 502–512 (2012).

5 Hoffmann, A., Levchenko, A., Scott, M. L. & Baltimore, D. The IκB–NF-κB signaling module: Temporal control and selective gene activation. Science 298, 1241–1245 (2002).

6 Lipniacki, T., Paszek, P., Brasier, A. R., Luxon, B. & Kimmel, M. Mathematical model of NF-κB regulatory module. J Theor Biol 228, 195–215 (2004).

7 Xaus, J. et al. LPS induces apoptosis in macrophages mostly through the autocrine production of TNF-α. Blood 95, 3823–3831 (2000).

8 Vandenbon, A., Teraguchi, S., Akira, S., Takeda, K. & Standley, D. M. Systems biology approaches to toll-like receptor signaling. WIREs Syst Biol Med 4, 497–507 (2012).

9 Zak, D. E., Tam, V. C. & Aderem, A. Systems-level analysis of innate immunity. Annu Rev Immunol 32, 547–577 (2014).

10 Werner, S. L., Barken, D. & Hoffmann, A. Stimulus specificity of gene expression programs determined by temporal control of IKK activity. Science 309, 1857–1861 (2005).

11 Purvis, J. E. & Lahav, G. Encoding and decoding cellular information through signaling dynamics. Cell 152, 945–956 (2013).

12 Caldwell, A. B., Cheng, Z., Vargas, J. D., Birnbaum, H. A. & Hoffmann, A. Network dynamics determine the autocrine and paracrine signaling functions of TNF. Genes Dev 28, 2120–2133 (2014).

13 Covert, M. W., Leung, T. H., Gaston, J. E. & Baltimore, D. Achieving stability of lipopolysaccharide-induced NF-κB activation. Science 309, 1854–1857 (2005).

14 Parameswaran, N. & Patial, S. Tumor necrosis factor-α signaling in macrophages. Crit Rev Eukaryot Gene Expr 20, 87–103 (2010).

15 Sung, M.-H. et al. Switching of the relative dominance between feedback mechanisms in lipopolysaccharide-induced NF-κB signaling. Sci Signal 7, ra6 (2014).

16 Murray, P. J. & Smale, S. T. Restraint of inflammatory signaling by interdependent strata of negative regulatory pathways. Nat Immunol 13, 916–924 (2012).

17 Kanarek, N., London, N., Schueler-Furman, O. & Ben-Neriah, Y. Ubiquitination and degradation of the inhibitors of NF-κB. Cold Spring Harb Perspect Biol 2 (2010).

18 Vereecke, L., Beyaert, R. & van Loo, G. The ubiquitin-editing enzyme A20 (TNFAIP3) is a central regulator of immunopathology. Trends Immunol 30, 383–391 (2009).

19 Hu, X., Chen, J., Wang, L. & Ivashkiv, L. B. Crosstalk among Jak-STAT, Toll-like receptor, and ITAM-dependent pathways in macrophage activation. J Leukoc Biol 82, 237–243 (2007).

20 Joo, J., Plimpton, S., Martin, S., Swiler, L. & Faulon, J.-L. Sensitivity analysis of a computational model of the IKK–NF-κB–IκB–A20 signal transduction network. Ann NY Acad Sci 1115, 220–239 (2007).

21 Basak, S., Behar, M. & Hoffmann, A. Lessons from mathematically modeling the NF-κB pathway. Immunol Rev 246, 221–238 (2012).

22 Williams, R. A., Timmis, J. & Qwarnstrom, E. E. Computational models of the NF-κB signalling pathway. Computation 2, 131–158 (2014).

23 Cheong, R., Hoffmann, A. & Levchenko, A. Understanding NF-κB signaling via mathematical modeling. Mol Syst Biol 4 (2008).

24 Sharp, G. C., Ma, H., Saunders, P. T. K. & Norman, J. E. A computational model of lipopolysaccharide-induced nuclear factor kappa B activation: a key signalling pathway in infection-induced preterm labour. PLoS One 8, e70180 (2013).

25 Pękalski, J. et al. Spontaneous NF-κB activation by autocrine TNFα signaling: a computational analysis. PLoS One 8, e78887 (2013).

26 Maiti, S., Dai, W., Alaniz, R. C., Hahn, J. & Jayaraman, A. Mathematical modeling of pro-and antiinflammatory signaling in macrophages. Processes 3, 1–18 (2015).

27 Lipniacki, T., Paszek, P., Brasier, A. R., Luxon, B. A. & Kimmel, M. Stochastic regulation in early immune response. Biophys J 90, 725–742 (2006).

28 Tay, S. et al. Single-cell NF-kB dynamics reveal digital activation and analogue information processing. Nature 466, 267–271 (2010).

29 Hughey, J. J., Gutschow, M. V., Bajar, B. T. & Covert, M. W. Single-cell variation leads to population invariance in NF-κB signaling dynamics. Mol Biol Cell 26, 583–590 (2015).

30 Lee, R. E. C., Walker, S. R., Savery, K., Frank, D. A. & Gaudet, S. Fold change of nuclear NF-κB determines TNF-induced transcription in single cells. Mol Cell 53, 867–879 (2014).

31 Hasenauer, J. et al. Identification of models of heterogeneous cell populations from population snapshot data. BMC Bioinform 12, 1–15 (2011).

32 Kalita, M. K. et al. Sources of cell-to-cell variability in canonical Nuclear Factor-κB (NF-κB) signaling pathway inferred from single cell dynamic images. J Biol Chem 286, 37741–37757 (2011).

33 Cheng, Z., Taylor, B., Ourthiague, D. R. & Hoffmann, A. Distinct single-cell signaling characteristics are conferred by the MyD88 and TRIF pathways during TLR4 activation. Sci Signal 8, ra69 (2015).

34 Paszek, P. et al. Population robustness arising from cellular heterogeneity. Proc Natl Acad Sci USA 107, 11644–11649 (2010).

35 Ravasi, T. et al. Generation of diversity in the innate immune system: macrophage heterogeneity arises from gene-autonomous transcriptional probability of individual inducible genes. J Immunol 168, 44–50 (2002).

36 Xue, J. et al. Transcriptome-based network analysis reveals a spectrum model of human macrophage activation. Immunity 40, 274–288 (2014).

37 Lu, Y. et al. Highly multiplexed profiling of single-cell effector functions reveals deep functional heterogeneity in response to pathogenic ligands. Proc Natl Acad Sci USA 112, E607–E615 (2015).

38 Altschuler, S. J. & Wu, L. F. Cellular heterogeneity: do differences make a difference? Cell 141, 559–563 (2010).

39 Satija, R. & Shalek, A. K. Heterogeneity in immune responses: from populations to single cells. Trends Immunol 35, 219–229 (2014).

40 Schuerwegh, A. J., Stevens, W. J., Bridts, C. H. & De Clerck, L. S. Evaluation of monensin and brefeldin A for flow cytometric determination of Interleukin-1 Beta, Interleukin-6, and Tumor Necrosis Factor-Alpha in monocytes. Comm Clin Cytometry 46, 172–176 (2001).

41 Rogers, S., Wells, R. & Rechsteiner, M. Amino acid sequences common to rapidly degraded proteins: the PEST hypothesis. Science 234, 364–368, doi:10.1126/science.2876518 (1986).

42 Merzlyak, E. M. et al. Bright monomeric red fluorescent protein with an extended fluorescence lifetime. Nat Methods 4, 555–557 (2007).

43 Rajasingh, J. et al. IL-10-induced TNF-alpha mRNA destabilization is mediated via IL-10 suppression of p38 MAP kinase activation and inhibition of HuR expression. FASEB J 20, E1393–E1403 (2006).

44 Brooks, S. A. & Blackshear, P. J. Tristetraprolin (TTP): interactions with mRNA and proteins, and current thoughts on mechanisms of action. Biochim Biophys Acta 1829, 666–679 (2013).

45 Carpenter, S., Ricci, E. P., Mercier, B. C., Moore, M. J. & Fitzgerland, K. A. Post-transcriptional regulation of gene expression in innate immunity. Nat Rev Immunol 14, 361–376 (2014).

46 Chuang, Y., Hung, M. E., Cangolese, B. K. & Leonard, J. N. Regulation of the IL-10-driven macrophage phenotype under incoherent stimuli. Innate Immun 22, 647–657 (2016).

47 Jayaraman, A. & Wood, T. K. Bacterial quorum sensing: signals, circuits, and implications for biofilms and disease. Annu Rev Biomed Eng 10, 145–167 (2008).

48 Schneider, C. A., Rasband, W. S. & Eliceiri, K. W. NIH Image to ImageJ: 25 years of image analysis. Nat Methods 9, 671–675 (2012).

49 Palumbo, M. C. et al. Collective behavior in gene regulation: post-transcriptional regulation and the temporal compartmentalization of cellular cycles. FEBS J 275, 2364–2371 (2008).

50 Aldridge, B. B., Burke, J. M., Lauffenburger, D. A. & Sorger, P. K. Physicochemical modelling of cell signalling pathways. Nat Cell Biol 8, 1195–1203 (2006).

51 Keogh, B. & Parker, A. E. Toll-like receptors as targets for immune disorders. Trends Pharm Sci 32, 435–443 (2011).

52 Gaba, A. et al. IL-10-mediated tristetraprolin induction is part of a feedback loop that controls macrophage STAT3 activation and cytokine production. J Immunol 189, 2089–2093 (2012).

53 Gais, P. et al. TRIF signaling stimulates translation of TNF-α mRNA via prolonged activation of MK2. J Immunol 184, 5842–5848 (2010).

54 Kafasla, P., Skliris, A. & Kontoyiannis, D. L. Post-transcriptional coordination of immunological responses by RNA-binding proteins. Nat Immunol 15, 492–502 (2014).

55 Bonizzi, G. & Karin, M. The two NF-κB activation pathways and their role in innate and adaptive immunity. Trends Immunol 25, 280–288 (2004).

56 Bode, J. G., Ehlting, C. & Häussinger, D. The macrophage response towards LPS and its control through the p38MAPK–STAT3 axis. Cell Signal 24, 1185–1194 (2012).

57 Radulescu, O., Gorban, A. N., Zinovyev, A. & Lilienbaum, A. Robust simplifications of multiscale biochemical networks. BMC Syst Biol 2 (2008).

58 Gutenkunst, R. N. et al. Universally sloppy parameter sensitivities in systems biology models. PLoS Comput Biol 3, 1871–1878 (2007).

59 Xue, Q. et al. Analysis of single-cell cytokine secretion reveals a role for paracrine signaling in coordinating macrophage responses to TLR4 stimulation. Sci Signal 8, ra59 (2015).

60 Raposo, G. & Stoorvogel, W. Extracellular vesicles: exosomes, microvesicles, and friends. J Cell Biol 200, 373–383 (2013).

61 Birtwistle, M. R. et al. Emergence of bimodal cell population responses from the interplay between analog single-cell signaling and protein expression noise. BMC Syst Biol 6 (2012).

62 To, T.-L. & Maheshri, N. Noise can induce bimodality in positive transcriptional feedback loops without bistability. Science 327, 1142–1145 (2010).

63 Busse, D. et al. Competing feedback loops shape IL-2 signaling between helper and regulatory T lymphocytes in cellular microenvironments. Proc Natl Acad Sci USA 107, 3058–3063 (2010).

64 Bagnall, J. et al. Quantitative analysis of competitive cytokine signaling predicts tissue thresholds for the propagation of macrophage activation. Sci Signal 11, eaaf3998 (2018).

65 Youk, H. & Lim, W. A. Secreting and sensing the same molecule allows cells to achieve versatile social behaviors. Science 343, 1242782 (2014).

66 Shalek, A. K. et al. Single-cell transcriptomics reveals bimodality in expression and splicing in immune cells. Nature 498, 236–240 (2013).

67 Shalek, A. K. et al. Single-cell RNA-seq reveals dynamic paracrine control of cellular variation. Nature 510, 363–369 (2014).

68 Postat, J., Olekhnovitch, R., Lemaître, F. & Bousso, P. A metabolism-based quorum sensing mechanism contributes to termination of inflammatory responses. Immunity 49, 654–665 (2018).

69 Bradley, J. R. TNF-mediated inflammatory disease. J Pathol 214, 149–160 (2008).

70 Peri, F. & Piazza, M. Therapeutic targeting of innate immunity with Toll-like receptor 4 (TLR4) antagonists. Biotechnol Adv 30, 251–260 (2012).

71 Chen, G. Y. & Nuñez, G. Sterile inflammation: sensing and reacting to damage. Nat Rev Immunol 10, 826–837 (2010).

72 Laskin, D. L. & Pendino, K. J. Macrophages and inflammatory mediators in tissue injury. Annu Rev Pharmacol Toxicol 35, 655–677 (1995).

73 Moore, K. J., Sheedy, F. J. & Fisher, E. A. Macrophages in atherosclerosis: a dynamic balance. Nat Rev Immunol 13, 709–721 (2013).

74 Parks, D. R., Roederer, M. & Moore, W. A. A new “Logicle” display method avoids deceptive effects of logarithmic scaling for low signals and compensated data. Cytometry A 69A, 541–551 (2006).

