## Supplementary Information for "Macrophages employ quorum licensing to regulate collective activation"

**This document includes:**

I. Experiments
- Figure S1
- Figure S2
- Table S1

II. Model
- Model formulation based upon prior knowledge
- Model formulation for other cellular processes
- Model development, parameterization, and implementation
- Figure S3
- Model parameterization: cell growth
- Model formulation: representing the experimental perturbations
- Variables, parameters, and equations

III. References

**Other supplementary materials include:**

- README.rtf
- Macrophage_model_homogeneous.m
- Macrophage_model_heterogeneous.m
- sim_cell.mat
- sim_population.mat
- Supplementary_tables.xlsx
- Flow_cytometry_data (folder)
- Confocal_microscopy_data (folder)

### I. Experiments

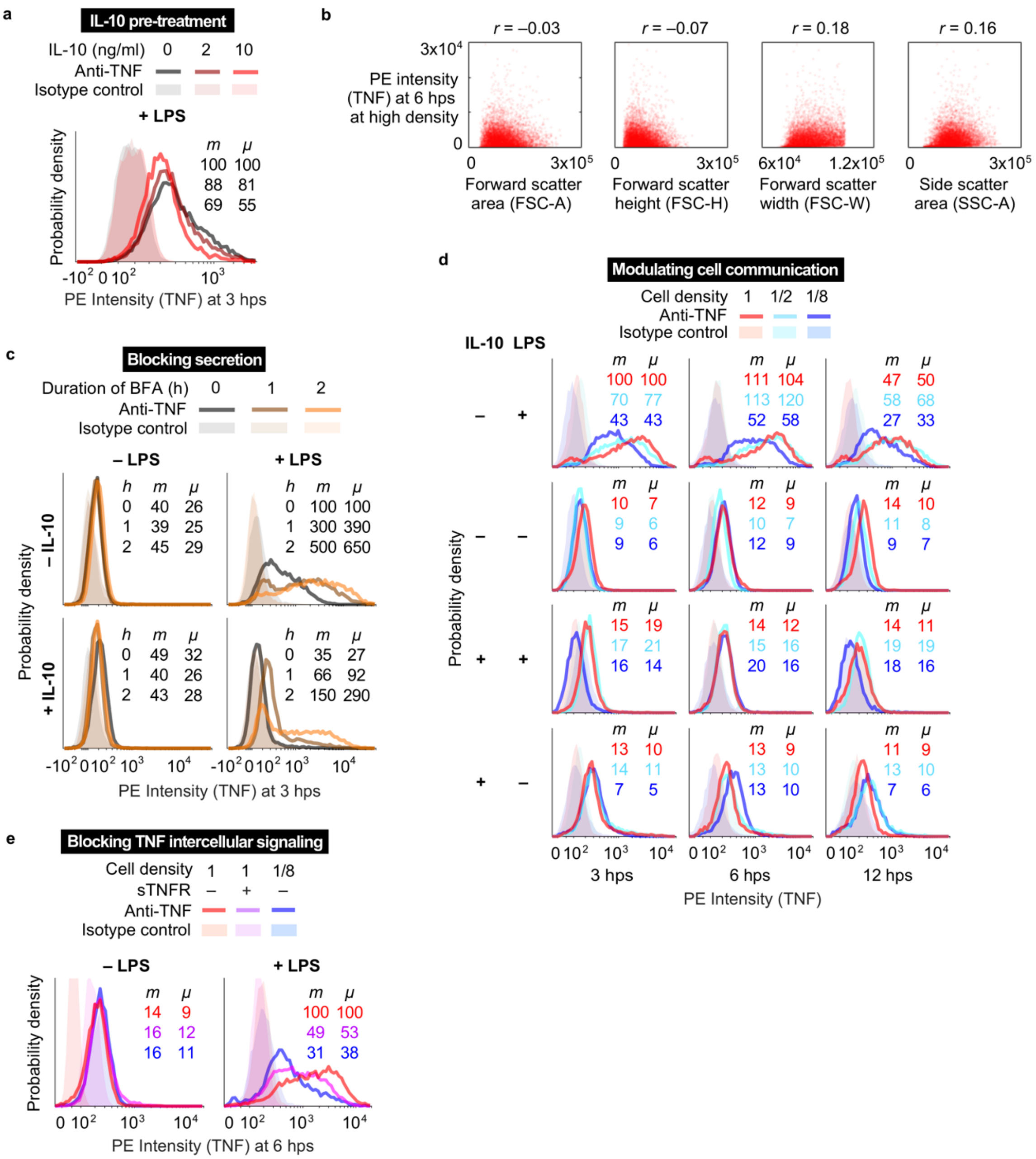


**Figure S1 | Heterogeneity in macrophage activation.** (**a**) Quantification of the effect of IL-10 pre-treatment from **Fig. 1b**. (**b**) TNF expression is generally uncorrelated with flow cytometric proxies for cell size. Axes are in linearly-scaled arbitrary units (a.u.). Low Pearson correlation coefficients (*r*) indicate little effect of cell size on TNF expression. (**c**) Quantification of the effect of BFA treatment from **Fig. 1c**. (**d**) Quantification of the effect of cell density from **Fig. 1d**. (**e**) Quantification of the effect of sTNFR pre-treatment from **Fig. 1e**. For each panel, median (*m*) and mean (*μ*) averages were calculated by subtracting the isotype control (shaded histograms) and normalizing to the condition assigned to a value of 100 for *m* and *μ*, *i.e.*, 100% of value for the reference case.

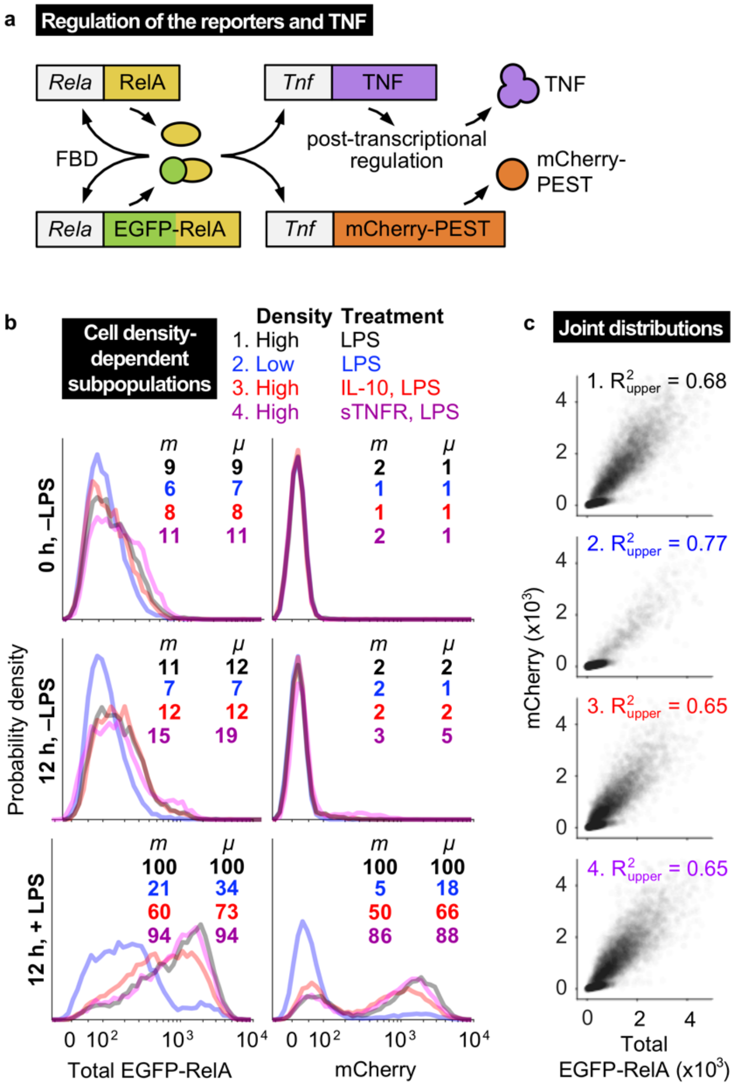


**Figure S2 | Single-cell and density-associated analysis of activation heterogeneity.** (**a**) Diagram of the regulation of endogenous RelA and TNF and the reporters EGFP-RelA and mCherry in reporter cells. (**b**) Quantification of reporter expression from **Fig. 2a**. Median (*m*) and mean (*μ*) averages were calculated by normalizing to the condition labeled 100%. X-axes are on a logicle scale. (**c**) Cells occupy high and low activation states. Data are from the four conditions at 12 hps in **b**. Within the cell subpopulation that undergoes high activation (subscripted as "upper"), reporters are highly correlated and similarly correlated across conditions. Axes are linearly scaled.

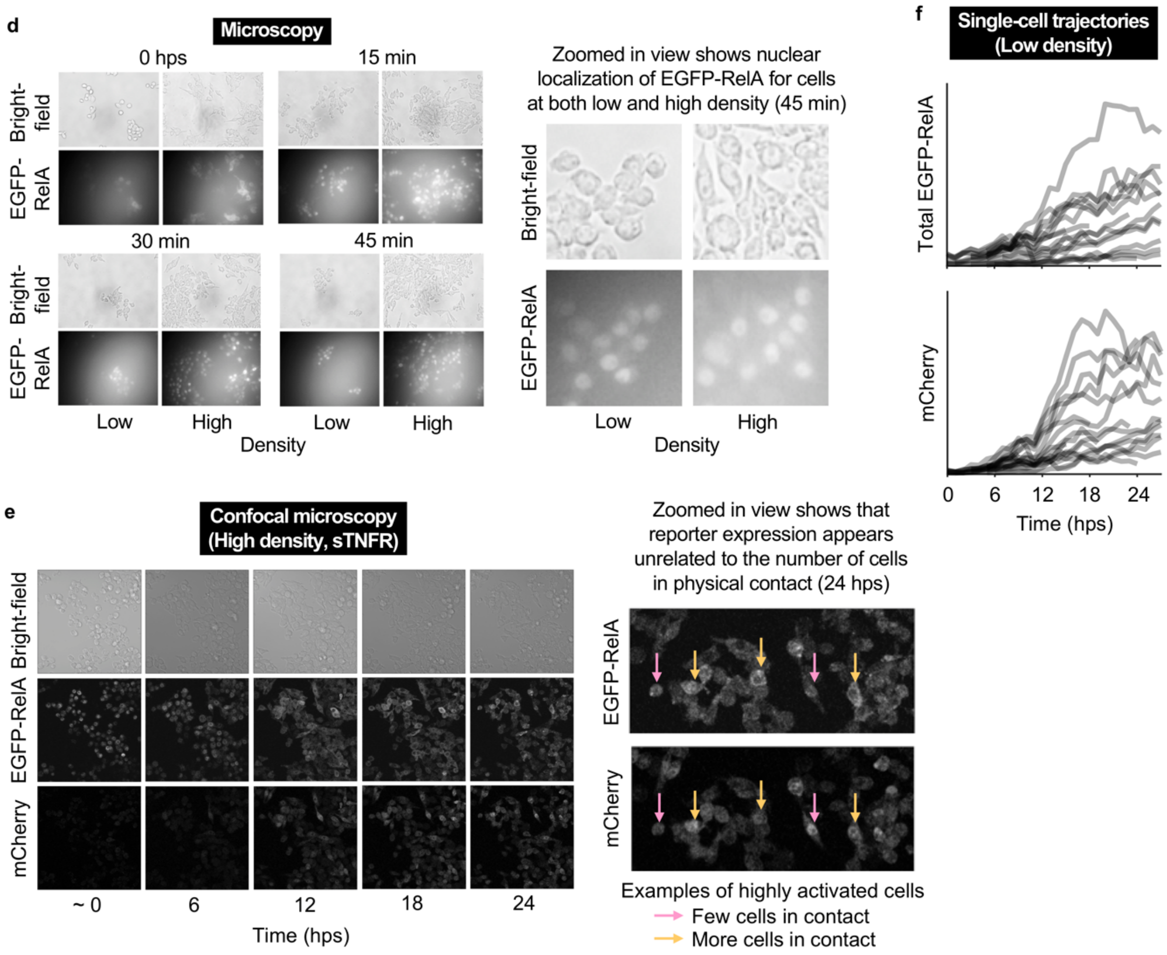


**Figure S2, continued.** (**d**) LPS induces EGFP-RelA nuclear translocation at both high and low cell density. Representative microscopy images show intracellular localization at several time points (*min* indicates minutes post-LPS stimulation). EGFP-RelA is initially primarily cytoplasmic, and after LPS treatment the nuclear intensity increases. Translocation occurs at both cell densities, indicating that low density does not prevent TLR4 signaling from activating NF-κB. (**e**) Direct cell-to-cell contact appears not to explain heterogeneous activation. Confocal microscopy images are shown for cells treated with sTNFR and LPS. "~0" indicates a timepoint shortly after LPS treatment. Cells with high fluorescence occur throughout the field of view, and some are adjacent to many cells while others are adjacent to few. Patterns were not apparent between a cell's fluorescence and the number or fluorescence intensity of neighboring cells. Although these snapshots do not capture the continuous history of cell-cell contact, as cells can migrate over time, the clumped cells show no discernible increase in fluorescence. For the purpose of visualization in **d**–**e**, images were uniformly brightened without altering the contrast. (**f**) Confocal microscopy trajectories for total EGFP-RelA and mCherry, showing 20 cells plated at low density. We note that some of the traces do not span the full duration of the timecourse due to mitosis or cells exiting the field of view.

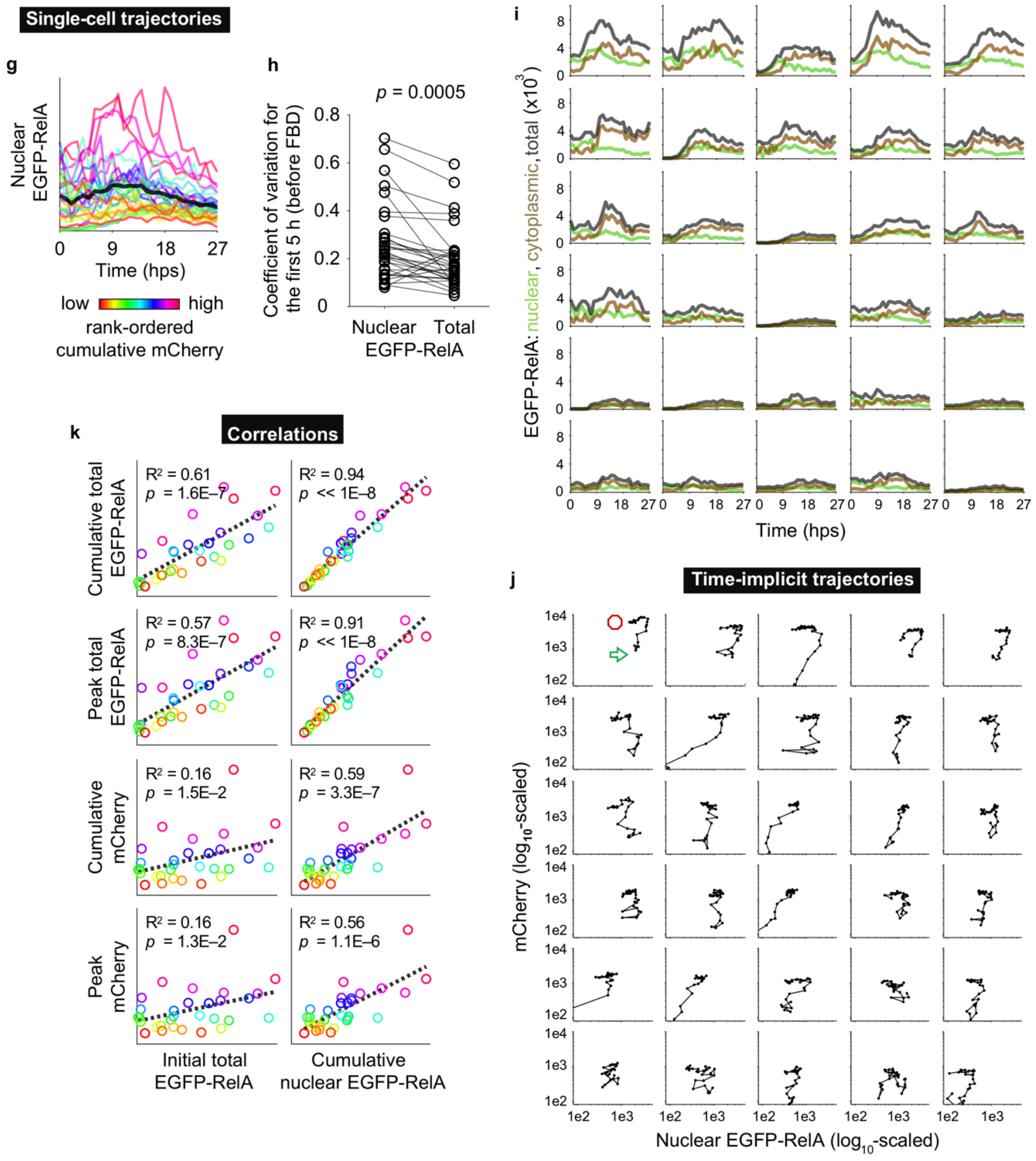


**Figure S2, continued.** (**g**–**k**) Analysis of reporter trajectories at high density with sTNFR and LPS. (**g**) Traces are color-coded by rank-ordered cumulative mCherry. (**h**) The coefficient of variation is greater for nuclear EGFP-RelA than total EGFP-RelA, consistent with nucleocytoplasmic translocations of the transcription factor. (**i**) Traces for nuclear (green), cytoplasmic (tan), and total (gray) EGFP-RelA are ordered row-wise from upper-left to lower-right by high-to-low cumulative mCherry. We observed that peak nuclear intensity generally preceded peak cytoplasmic intensity. Axes are linearly scaled. (**j**) Time-implicit trajectories of nuclear EGFP-RelA and mCherry vary in activation magnitude but follow a characteristic pattern—starting in the lower-left, moving to the upper-right, and then moving left to the end point. The green arrow and red octagon in the upper-left panel indicate the start and finish of the timecourse. Axes are $\log_{10}$-scaled. (**k**) Initial EGFP-RelA and integrated nuclear EGFP-RelA are predictive for total EGFP-RelA and mCherry. *P*-values indicate significance from a one-tailed test for the Pearson correlation.

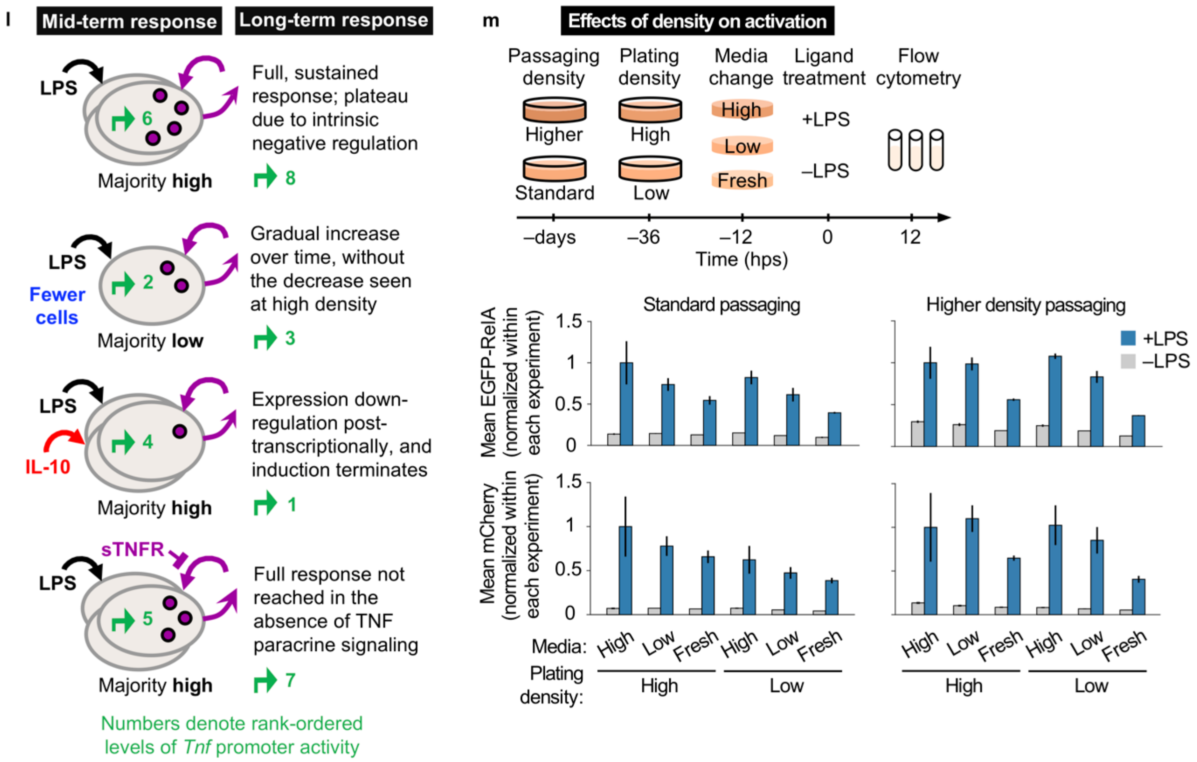


**Figure S2, continued.** (**l**) The diagram provides a conceptual summary of observations on *Tnf* promoter activity (green arrow) and TNF protein expression (purple circles) for mid-term (12 hps) and long-term (24 hps) responses across the experimental perturbations. With microscopy, the range of values is narrower than with flow cytometry. Additionally, the readout in fluorescence a.u. is specific to each experiment, such that typically the most comparable features are those that are magnitude-independent, *e.g.*, trajectory shape and the time to reach peak intensity. To enable more comparisons, we used the mean average values in **Fig. 2a** (in comparable flow cytometry a.u.) to scale the microscopy data in **Fig. 2f** at 12 hps to comparable a.u. from which long-term responses could also be estimated. This analysis was used to rank *Tnf* promoter activity states (based on normalized mCherry expression) from low (1) to high (8) (green text). (**m**) Modulating intercellular communication through the cell density during passaging, at plating, and for the media change. Data for standard passaging and higher density passaging were collected separately and are in distinct a.u. Data are normalized within each fluorescent reporter readout. The line with each bar indicates +/– one standard deviation for the mean of three biologic replicates.

**Table S1. Hypothesis testing of cell density effects[1]**

Hypothesis test A

| | | Standard passaging | | | | Higher density passaging | | | |
|---|---|---|---|---|---|---|---|---|---|
| | | High density | | Low density | | High density | | Low density | |
| | **Media** | NT | LPS | NT | LPS | NT | LPS | NT | LPS |
| % High | High vs. Low | 5.0E-01 | 8.3E-02 | 3.2E-04 | 7.9E-03 | 6.1E-03 | 5.0E-01 | 3.9E-04 | 1.2E-02 |
| | High vs. Fresh | 3.6E-01 | 4.4E-02 | 9.7E-05 | 5.5E-04 | 4.1E-04 | 3.8E-03 | 2.1E-05 | 1.7E-05 |
| | Low vs. Fresh | 7.3E-03 | 2.0E-01 | 3.3E-04 | 1.5E-03 | 3.8E-03 | 4.5E-04 | 3.2E-05 | 4.2E-05 |
| Mean | High vs. Low | 5.0E-01 | 8.5E-02 | 3.7E-05 | 1.8E-02 | 2.8E-02 | 4.6E-01 | 7.2E-04 | 2.7E-03 |
| | High vs. Fresh | 7.7E-02 | 2.1E-02 | 1.8E-04 | 4.4E-04 | 1.2E-04 | 1.1E-02 | 5.0E-05 | <1E-05 |
| | Low vs. Fresh | 8.9E-04 | 1.2E-02 | 4.9E-03 | 5.0E-03 | 9.1E-04 | 3.6E-04 | <1E-05 | 1.8E-04 |

Hypothesis test B

| | | Standard passaging | | | | | | Higher density passaging | | | | | |
|---|---|---|---|---|---|---|---|---|---|---|---|---|---|
| | | High | | Low | | Fresh | | High | | Low | | Fresh | |
| | **Cell density** | NT | LPS | NT | LPS | NT | LPS | NT | LPS | NT | LPS | NT | LPS |
| % High | High vs. Low | 5.0E-01 | 2.5E-01 | 6.3E-05 | 1.9E-02 | 7.5E-05 | 2.4E-04 | 2.6E-03 | 3.8E-01 | 2.4E-04 | 2.3E-03 | 1.6E-05 | 1.5E-05 |
| Mean | High vs. Low | 5.0E-01 | 1.6E-01 | 2.4E-04 | 6.5E-02 | 1.3E-03 | 4.0E-03 | 7.8E-03 | 5.0E-01 | 8.6E-04 | 3.2E-02 | <1E-05 | 1.3E-05 |

Hypothesis test C

| | | High density | | | | | | Low density | | | | | |
|---|---|---|---|---|---|---|---|---|---|---|---|---|---|
| | | High | | Low | | Fresh | | High | | Low | | Fresh | |
| | **Passaging** | NT | LPS | NT | LPS | NT | LPS | NT | LPS | NT | LPS | NT | LPS |
| % High | High vs. stand. | 1.2E-05 | 8.1E-03 | <1E-05 | 2.9E-05 | <1E-05 | 1.6E-05 | <1E-05 | 1.7E-04 | <1E-05 | 1.5E-05 | <1E-05 | <1E-05 |

1. *P*-values were calculated from one-tailed *t*-tests for the effects of various conditions on EGFP-RelA expression (**Supplementary Fig. S2m**). Tests were conducted using the mean EGFP-RelA expression and the percentage of cells above a threshold value. Green highlighting indicates statistical significance ($\alpha < 0.05$) following a Bonferroni correction for multiple hypothesis testing (dividing $\alpha$ by the number of hypotheses $m$). In test A, the null hypothesis is that media change has no effect, and the alternative hypothesis is that media from cells plated at high density leads to more EGFP-RelA than media from cells plated at low density, which leads to more EGFP-RelA than with fresh media ($m$ = 12). In test B, the null hypothesis is that cell density has no effect, and the alternative hypothesis is that cells plated at high density have more EGFP-RelA than cells plated at low density ($m$ = 6). In test C, the null hypothesis is that the density during passaging has no effect, and the alternative hypothesis is that cells passaged at higher density have more EGFP-RelA than cells passaged at standard density ($m$ = 6). Among the three tests, the most consistently significant effects are in C.

### II. Model

We developed a computational model to incorporate the new findings with prior knowledge on macrophage activation. We focused the scope to processes to which the new findings most directly pertain. This section describes prior knowledge and the model formulation, development, parameterization, and analysis. Matlab files for homogeneous and heterogeneous models are provided as supplementary materials, available at github.com/bagherilab, and additionally detailed in **Supplementary Tables S2–S5**.

#### Model formulation based upon prior knowledge

For each stage of the TNF response to LPS, known cellular mechanisms are described followed by how they are represented in the model. Salient state variables are specified in parentheses.

***TLR4 signaling*** ($\dot{x}_1$, $\dot{x}_2$, $\dot{x}_5$, $\dot{x}_6$, $\dot{x}_{EC2}$)

*Mechanism:* LPS activates TLR4 signaling through two pathways named after their use the adaptor proteins MyD88 (Myeloid differentiation primary response gene 88) and TRIF (Toll/interleukin-1 receptor (TIR)-domain-containing adapter-inducing interferon-β)[1]. While these pathways differ in certain components and downstream targets, they overlap in NF-κB activation. MyD88 signaling involves the formation of a multi-subunit protein complex at the plasma membrane called the Myddosome, whose activity is induced and terminated rapidly. TRIF signaling requires TLR4 internalization to endosomes; it is initially delayed as activated TLR4 begins to reversibly shuttle from the plasma membrane to endosomes, but it is longer-lasting than MyD88 signaling. Signaling from activated receptors terminates upon maturation of early endosomes to late endosomes.

*Model:* LPS has an initial value of 1 a.u., corresponding to a dose of 100 ng/ml. The inactive (TLR4) and active (TLR4*) forms of the receptor are assigned initial values of 0.1 and 0 a.u., respectively. Since LPS is in large molar excess of the receptor, its loss over time can be represented simply by first-order degradation. TLR4 is synthesized constitutively in its inactive form and undergoes first-order degradation. Receptor activation depends on LPS dose, not cell density. For the signaling cascade, we developed a reduced complexity version of a previously published model[1], *e.g.*, that does not distinguish MyD88-mediated and TRIF-mediated signaling. As a general principle, we used a minimal number of variables to represent the most salient processes. During model development, we observed that reducing the granularity of certain mechanisms had little impact on the dynamics of interest such as NF-κB translocation.

***NF-κB activation*** ($\dot{x}_7$, $\dot{x}_8$, $\dot{x}_{10}$, $\dot{x}_{11}$, $\dot{x}_{13}$, $\dot{x}_{14}$, $\dot{x}_{15}$, $\dot{x}_{16}$, $\dot{x}_{17}$, $\dot{x}_{18}$)

*Mechanism:* NF-κB is a dimer composed from five subunits: RelA (p65), cRel, RelB, p50, and p52[2]. In TLR4 signaling and in TNFR signaling, IKKK phosphorylates IKK (IκB kinase), which phosphorylates IκB, targeting IκB for degradation. The de-sequestered NF-κB then translocates to the nucleus and induces target gene transcription. NF-κB is eventually re-sequestered by IκB, and NF-κB–IκB translocates to the cytoplasm. This activation and inactivation comprise one cycle of NF-κB nucleocytoplasmic translocation[1].

*Model:* we reduced and modified a portion of a previous model[1] while retaining NF-κB oscillatory behavior. Since RAW reporter cells express a functional EGFP-p65, and p65/p50 is the primary NF-κB dimer, we represent EGFP-p65-containing and native p65/p50 dimers as the same NF-κB variable. The variable IκB represents the IκBα gene product[3].

***RelA feedback dominance switching*** ($\dot{x}_9$, $\dot{x}_{11}$)

*Mechanism:* RAW cells treated with LPS above a certain dose threshold enter a positive feedback loop in which NF-κB induces *Rela* expression[4]. This feedback dominance (FBD) switch overpowers the negative feedback from the induction of IκB by NF-κB. Since the switch requires *de novo* expression of the TF Ikaros, it takes effect starting several hours post-LPS. LPS at 100 ng/ml is well above the dose threshold.

*Model:* NF-κB undergoes the FBD switch in all cells at and above a presumed dose of 1 ng/ml. A time-dependent function was formulated for NF-κB-induced activity at the *Rela* promoter, based on previously published timecourse ChIP data[4] for NF-κB localization at the *Rela* promoter in RAW cells.

***Early regulation of Tnf translation*** ($\dot{x}_{24}$, $\dot{x}_{25}$)

*Mechanism:* in the resting cell state, *Tnf* mRNA lacks a poly(A) tail and is not translated[5]. After LPS treatment, the mRNA is polyadenylated, allowing poly(A)-binding protein (PABP) to pseudo-circularize the mRNA, which increases ribosome recycling for rapid translation. TRIF signaling also promotes translation, by activating p38 mitogen activated protein kinase (MAPK), which activates MAP kinase-activated protein kinase 2 (MK2) to phosphorylate eukaryotic translation initiation factor 4E (eIF4E), which binds to the 5' mRNA cap and recruits the 40S ribosomal subunit. TRIF signaling also dephosphorylates eIF2, which de-represses translation by recruiting the 60S subunit[5].

*Model:* Since TNF production requires LPS treatment, the initial values of *Tnf* mRNA and TNF protein are set to zero regardless of the initial value of NF-κB.

***Tnf post-transcriptional regulation*** ($\dot{x}_{19}$, $\dot{x}_{20}$, $\dot{x}_{24}$, $\dot{x}_{25}$)

*Mechanism: Tnf* mRNA is regulated post-transcriptionally through AU-rich elements (AREs) in its 5' UTR, with binding sites for over 20 proteins[5]. Some proteins such as Tristetraprolin (TTP) destabilize the mRNA by recruiting deadenylases and degradation factors, and others such as Hu-antigen R (HUR) stabilize the mRNA by competing with destabilizing proteins for occupancy[5]. In unstimulated macrophages, TTP is expressed at low levels[6]. Shortly after LPS treatment, kinases including p38 and Erk are activated and *Tnf* mRNA is stabilized[7]. However, TLR4 and TNFR signaling also induce TTP expression via p38 and ERK signaling[8] (and IL-10R signaling also induces TTP expression, via STAT3[6]). TTP binds to *Tnf* mRNA and leads to its destabilization[9]. The outcome is a limited-duration burst in TNF expression[7].

*Model:* To capture the limited-duration burst in TNF expression while limiting model complexity, stabilizing regulation (SR) is represented by one variable and destabilizing regulation (DSR) is represented by another. SR becomes active after LPS (downstream of TLR4* and TNFR* via IKKK*), and it decreases in activity over time. SR slows the degradation of *Tnf* mRNA. DSR becomes active after a delay. DSR increases *Tnf* mRNA degradation and suppresses TNF translation.

***Action of IL-10 pre-treatment through STAT3*** (represented through a parameter, not state variables)

*Mechanism:* Bcl-3 is a nuclear protein that dimerizes with p50 or p52 and binds NF-κB-responsive promoters[10]. Bcl-3 expression is induced by STAT3 in IL-10R signaling.

*Model:* To limit model complexity, rather than introducing variables for IL-10R signaling, STAT3 activation, or Bcl-3 interactions, we found that representing the effect of IL-10 on FBD via a fitted parameter was sufficient to capture the observed decrease in reporter expression (**Fig. 2a**).

***Action of IL-10 pre-treatment through MAPKs*** ($\dot{x}_{19}$, $\dot{x}_{20}$)

*Mechanism:* MAPKs regulate the initial response to LPS and the resolution[11]. Their effects on downstream targets are complex and have been described as *incoherent*, *i.e.*, having seemingly opposing effects. Signaling via MyD88 and TRIF activates p38, extracellular-signal-regulated kinases 1 and 2 (ERK1/2), and c-Jun N terminal kinase (JNK). MK2, a phosphorylation target of p38, regulates *Tnf* and *Il10* mRNA stability by: (a) preventing recruitment of the adenylase CCR4-associated factor 1 (CAF1), thereby preventing proteins like TTP from destabilizing target mRNAs, and (b) inducing TTP expression. IL-10R signaling also regulates *Tnf*, by activating STAT3 (which induces TTP) and increasing expression of dual specific phosphatase 1 (DUSP1, which dephosphorylates p38 and inhibits late-phase p38 activity)[11]. In macrophages, IL-10R signaling destabilizes inflammatory cytokine mRNAs like *Tnf* that contain 3' UTR AU-rich elements (AREs), by: (a) repressing LPS-induced activation of p38 MAPK, and (b) inhibiting expression of HuR, a protein that stabilizes mRNAs by binding AREs[12].

*Model:* IL-10 treatment decreases stabilizing regulation and increase destabilizing regulation of *Tnf* mRNA. Effects of the MAPKs are represented by SR downstream of IKKK activation, and effects of TTP are represented by DSR. IL-10 pre-treatment prevents SR, and it activates DSR by 0 hps.

***TNFR activation*** ($\dot{x}_{3}$, $\dot{x}_{4}$, $\dot{x}_{5}$, $\dot{x}_{6}$, $\dot{x}_{EC1}$)

*Mechanism:* Extracellular TNF binds TNFR1, and the receptor-ligand complex is internalized[13]. Adaptor proteins activate IKKK and MAPK signaling, leading to NF-kB activation.

*Model:* the TNF pool has an initial value of 0 a.u. The inactive (TNFR) and active (TNFR*) forms of the receptor are assigned initial values of 0.1 and 0 a.u., respectively, analogous to the TLR4 receptor. TNFR* and TLR4* converge at IKKK activation, and thus overlap in regulating *Tnf*.

***Cell density*** ($\dot{x}_{EC1}$)

*Mechanism:* Cell density affects the proportion of highly activated cells and the amount of secreted TNF.

*Model:* To account for population growth over time, the rate at which secreted TNF contributes to the extracellular TNF pool is multiplied by a time-dependent function. The multiplier is for the pool, not individual cells. The proportion of highly activated cells is based on experimental observation.

***Blockade of TNFR signaling*** ($\dot{x}_3$, $\dot{x}_4$)

*Mechanism:* At 1 h pre-LPS, cells were treated with soluble TNF receptor (sTNFR), which competes with surface TNFR for TNF. The dose of sTNFR was chosen based on a prior study such that it would be in molar excess of secreted TNF[14].

*Model:* sTNFR blockade prevents TNFR activation. As a result, following LPS treatment, downstream nodes such as IKKK are activated to a lesser extent than they would be with paracrine feedback.

***Secretion*** ($\dot{x}_{25}$, $\dot{x}_{EC1}$)

*Mechanism:* p38 modulates the activity of proteins involved in endocytic trafficking to enhance cytokine secretion during the LPS response. At the cell surface, TNF precursor is released in soluble form following cleavage by TNFα-converting enzyme (TACE), which is activated by an LPS-activated lipid hydrolase[11].

*Model:* since simulations begin at 0 hps, TNF secretion is assigned a constant rate parameter.

***Blockade of secretion*** ($\dot{x}_{25}$, $\dot{x}_{EC1}$)

*Mechanism:* Brefeldin A (BFA) prevents secretion involving Golgi transport. After BFA treatment, TNF is no longer secreted, and it accumulates intracellularly.

*Model:* A time-dependent step-down function represents prevention of TNF secretion. Since some TNF is secreted prior to BFA, some paracrine signaling occurs, though to a lesser extent than without BFA.

### Model formulation for other cellular processes

***Receptor activation***

Receptors are synthesized constitutively in an inactive form and are initially at steady state. Receptors become activated through a second-order reaction with the extracellular cue, and initiate the downstream cascade. Certain rate constants differ for TLR4 and TNFR.

***Kinase cascade***

Kinases are present at a constant level. They are initially inactive, and become activated by a second-order reaction with an upstream node. Basal deactivation is first-order. For example, TLR4* and IKKK react to convert IKKK to IKKK*, which eventually returns to IKKK (due to the action of phosphatases).

***Translocation***

Translocation between the cytoplasm and nucleus is first order. Rate constants differ by species (NF-κB, IκB, and NF-κB–IκB) and direction of movement.

***Transcription***

Transcription was formulated using fractional activation. Terms for transcriptional activators are in both the numerator and denominator. Terms for inhibitory effects are in the denominator.

***Translation***

Translation is generally treated as first order with mRNA. However, *Tnf* translation has additional regulation.

***Secretion***

There are 30 cells in the model, and the sum of their TNF secretion contributes to the extracellular pool.

***Degradation***

Degradation is generally treated as first order. Cases involving regulated degradation are nonlinear.

### Model development, parameterization, and implementation

Before parameterizing the full model, we started with a section (cell-intrinsic NF-κB module) that includes TLR4 signaling, regulation of NF-κB activation, and RelA and IκB expression (**Supplementary Fig. S3a**). Following LPS treatment, TLR4 is activated to TLR4*, which activates IKKK to IKKK*, which activates IKK to IKK*. In the cytoplasm, IKK* associates with IκB or NF-κB–IκB to form IKK–IκB or NF-κB–IKK–IκB, respectively. IKK–IκB and NF-κB also associate to form NF-κB–IKK–IκB, and IKK targets IκB for degradation. Formally, degradation occurs regardless of whether NF-κB is complexed, but for the purpose of model reduction, we represented this process for NF-κB–IKK–IκB only; we observed that this decision did not have a noticeable impact on simulations. The reaction releases NF-κB and IKK. NF-κB can then enter the nucleus and induce transcription of *Rela* and *Ikba*. NF-κB, IκB, and NF-κB–IκB translocate between the nucleus and cytoplasm. Since NF-κB–IκB exits the nucleus much faster than it enters, its translocation is treated as unidirectional; as with the representation of IKK's effect on IκB, this model reduction had no discernable impact. During model development, we encountered many such instances where complexity could be reduced, such as for reasons relating to separation of timescales, redundant pathway effects, or negligible reaction fluxes.

To conduct an unbiased search of parameter space for fitting the model, we tested many parameter sets using a Sobol sequence—a pseudorandom number list that uniformly samples the unit hypercube in the limit of the sequence[15]. This initial sweep was followed by multi-objective optimization using a genetic algorithm with multiple generations comprising the following steps:

- Evaluate: quantify goodness of fit for each parameter set based on the deviation of simulated outcomes from experimental data.
- Select: identify parameter sets that yield the best fit within the current population. We applied "all-or-nothing" criteria for whether population-mean simulated outcomes fell within specified windows of acceptable deviation from population-mean experimental data.
- Repopulate: copy the selected sets and restore the original population size.
- Mutate: introduce random variation to the parameter values, to generate variation on which the above operations can be applied in subsequent generations. Variation was drawn from Gaussian distributions centered on current-generation estimated values, with a simulated annealing approach of decreasing the coefficients of variation as the number of generations increased.

The genetic algorithm yielded a family of similar-performing four-parameter sets for the NF-κB module that were carried forward and sampled during the fitting of the remaining eight parameters in the full model as described in **Results** (**Fig. 3**). The calibration resulted in a one-cell (*i.e.*, homogeneous) model.

Heterogeneity was then incorporated by creating a population of 30 cells—each with 25 intracellular state variables with the same reaction terms—and assigning different basal transcription rates to the RNA for NF-κB, and different initial values to the state variables for the RNA for NF-κB and for cytoplasmic NF-κB–IκB. These values are proportional to fluorescence intensity measurements (in a.u.) of the 30 cells at high density quantified from confocal microscopy. Values for cells at low density were obtained in equivalent units as described in the **Results** for **Fig. 3b**. Fitting, simulations, analysis, and visualization steps were conducted in Matlab.

To highlight the NF-κB module independent of TNF paracrine feedback, representative simulations are included for the homogeneous model under varied conditions (**Supplementary Fig. S3b**). The simulations show that following TLR4 activation and signaling, NF-κB is activated to enter the nucleus and induce target gene transcription. Damped oscillations occur due to negative feedback between NF-κB activation and IκB expression and the time required for *de novo* IκB expression. Depending on the LPS dose, FBD switching, and initial value of cytoplasmic inactive NF-κB, qualitatively distinct trajectories for total and nuclear NF-κB are produced, consistent with the wide-ranging outcomes observed in confocal microscopy (**Fig. 2, Supplementary Fig. S2**).

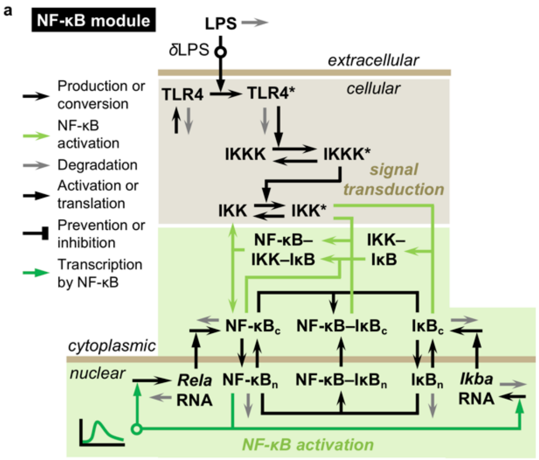


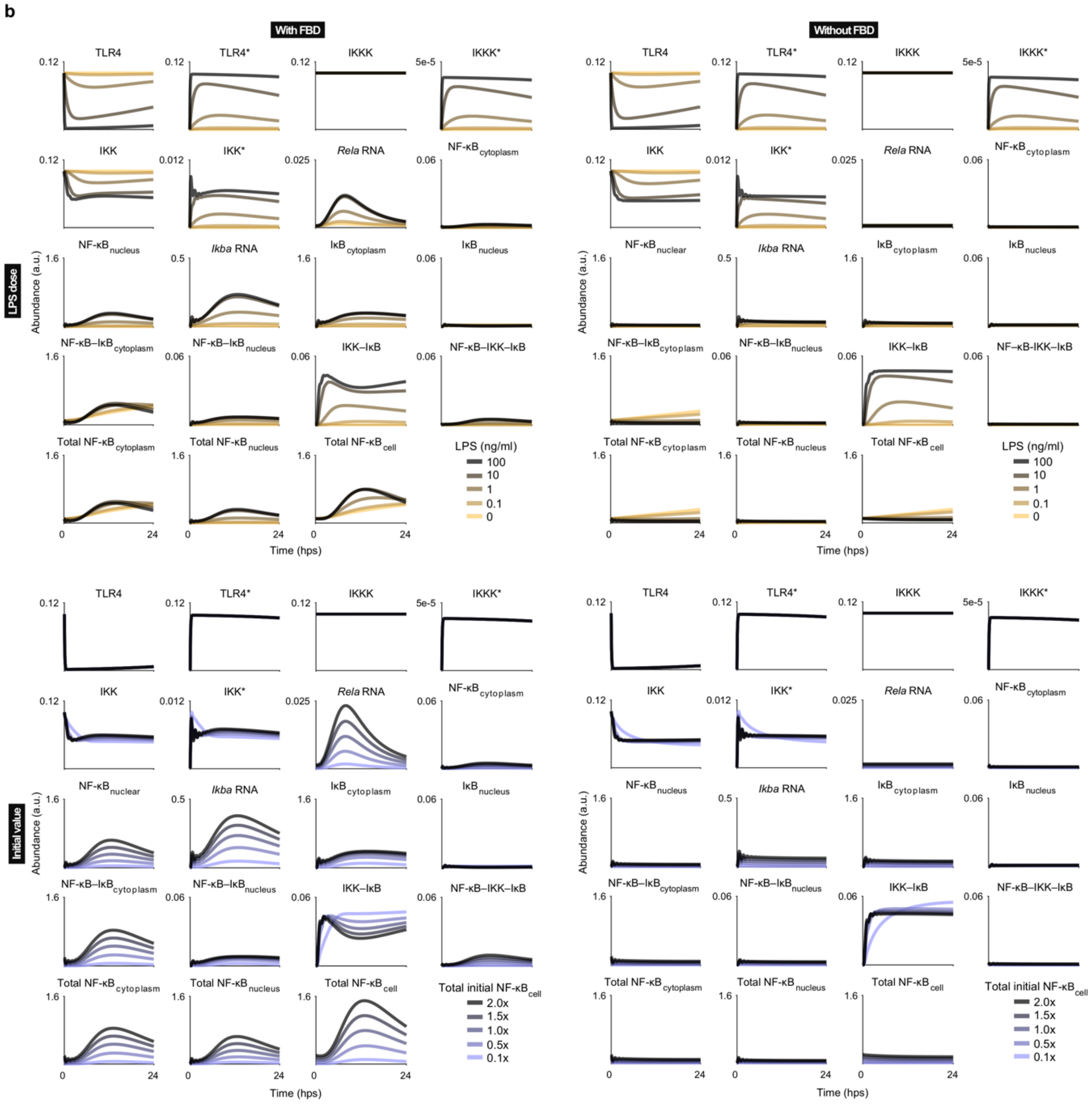


**Figure S3 | Computational model development** (**a**) Diagram of the NF-κB module. Arrows denote the processes indicated in the legend. State variables are in bold non-italicized text. (**b**) Simulations across LPS doses and NF-κB initial values (in the inactive cytoplasmic fraction), with and without the FBD switch. The base case values are 100 ng/ml dose and *1x* initial value (0.1 a.u.). The first 16 panels are individual state variables, and the last three panels are the total cytoplasmic, total nuclear, and total-cell amounts of NF-κB determined by summing the individual variables.

### Model parameterization: cell growth

RAW cell density was monitored across time points and conditions. These data were used to fit a logistic model for time-dependent growth for high and low density plating (**Supplementary Fig. S3c**).

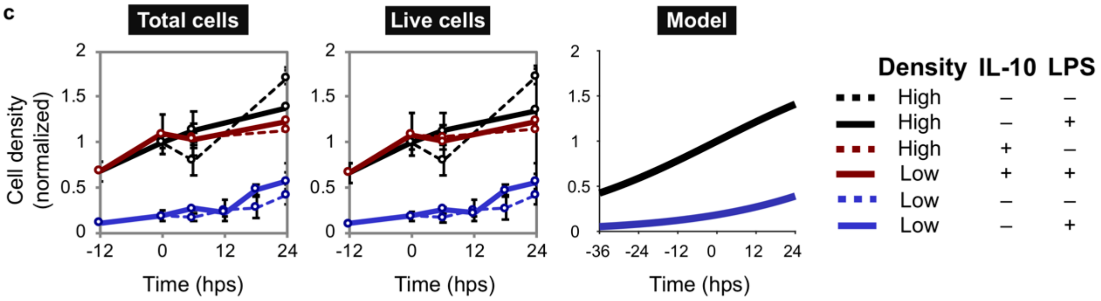


**Figure S3, continued.** (**c**) Population growth. RAW cells were plated at high (3.3x10[5] cells/ml) or low (4.1x10[4] cells/ml) density (at –36 hps). To quantify cell density from the same plates over time, representative images were obtained by bright-field microscopy at several time points for automated analysis in ImageJ. Circles are mean averages, and error bars are +/– one standard deviation as determined from five images from each of two replicate plates. Y-axis units are normalized such that high density at 0 h is defined as 1 a.u. *Left:* growth was unaffected by IL-10 pre-treatment and LPS treatment. *Middle:* cell viability as estimated from Trypan blue staining remained ~100% across conditions. *Right:* data were used to fit a model for cell growth as a function of time and the density at plating, described below.

### Model formulation: representing the experimental perturbations

Experimental perturbations were modeled using the equations below and as depicted in the diagram (**Supplementary Fig. S3d**). Parameters are denoted by $k$ for kinetic processes and $w$ for weights in transcriptional regulation. Variables are abbreviated as: NFkBn, nuclear NF-κB; Tnfm, *Tnf* mRNA; TNF, intracellular TNF; TNFpool, extracellular TNF pool; SR, stabilizing regulation on *Tnf* mRNA; DSR, destabilizing regulation on *Tnf* mRNA.

The time-dependent cell density ($\rho$) is described by a logistic equation. Units are relative to high cell density (1 a.u.) at the time of LPS treatment ($t$ = 0). From a fit to data (**Supplementary Fig. S3a**), and using constraints for (1) eight-fold difference in plating and (2) equal density in the limit of time, the horizontal asymptote for maximum density ($r_1$) is 1.99 a.u. and the rate of logistic growth ($r_2$) is 0.0365 $h^{-1}$ for high or low density populations. $\tau_{\text{density}}$ is –0.247 h for high density and 62.6 h for low density.

$$\rho(t, \tau_{\text{density}}) = \frac{r_1}{1 + e^{-r_2 \cdot (t - \tau_{\text{density}})}}$$

$$\text{Flux of TNF secreted by the cell population} = \rho \cdot (1 - \delta_{\text{sTNFR}}) \cdot k_{\text{secretion}} \cdot \sum_i^N [\text{TNF}_i]$$

Terms with $\delta$ indicate the presence or absence of a perturbation:

$$\delta_{\text{LPS}} = \begin{cases} no\ \text{LPS} \longrightarrow 0 \\ \text{LPS} \longrightarrow 1 \end{cases}$$

$$\delta_{\text{IL10}} = \begin{cases} no\ \text{IL10} \longrightarrow 0 \\ \text{IL10} \longrightarrow 1 \end{cases}$$

Transcription of *Rela*: the feedback dominance switch is represented by a time-dependent function ($F_{FBD}$) multiplied by the term for NF-κB positive feedback. $F_{FBD}$ was formulated and fitted based on timecourse ChIP data[4] for RelA localization at the *Rela* promoter post-LPS.

$$F_{FBD} = \frac{1}{6}\delta_{LPS}\delta_{FBDLic}t^3e^{-t}$$

$$\text{Flux of } Rela \text{ transcription} = k_{tx}\frac{w_{basaltxNFkB} + F_{FBD}w_{NFkBtxNFkB}[NFkBn]}{1 + F_{FBD}w_{NFkBtxNFkB}([NFkBn] + w_{IL10txFBD}\delta_{IL10})}$$

Transcription of *Tnf* and *mCherry*: nuclear NF-κB induces transcription at the *Tnf* promoter.

$$F_{NFkB\_Tnf} = w_{NFkBtxTnf}[NFkBn]$$

With IL-10 pre-treatment, there is a decrease in LPS-induced transcription at the *Tnf* promoter that begins after some delay. This is described non-mechanistically by a time-dependent ramp-down ($F_{IL10\_Tnf}$) starting at time $\tau_{IL10\_1}$ and ending at time $\tau_{IL10\_2}$, where $H$ is the Heaviside function.

$$F_{IL10\_Tnf} = 1 - \delta_{IL10}\left(1 - \frac{\max(t - \tau_{IL10\_1}, 0)}{\tau_{IL10\_2} - \tau_{IL10\_1}}\right)\left(1 - \frac{1}{1 + H(t - \tau_{IL10\_2})}\right)$$

$$\text{Flux of transcription from the } Tnf \text{ promoter} = \delta_{LPS}\delta_{TnfLic}k_{tx}\frac{F_{NFkB\_Tnf}}{1 + F_{NFkB\_Tnf}}F_{IL10\_Tnf}$$

Post-transcriptional and translational regulation of *Tnf*: stabilizing regulation slows the degradation of *Tnf* mRNA, and destabilizing regulation promotes degradation. These effects act in opposition.

$$F_{stabilize} = \frac{1}{1 + w_{stabilize}[SR]}$$

$$F_{destabilize} = \frac{w_{destabilize}[DSR](w_{maxdestabilize} - 1)}{1 + w_{destabilize}[DSR]}$$

$$\text{Flux of } Tnf \text{ mRNA degradation} = k_{degTnfm}[Tnfm](F_{stabilize} + F_{destabilize})$$

Destabilizing regulation also represses the translation of *Tnf*.

$$F_{repress\_tl} = \frac{1 + w_{destabilize}[DSR]}{1 + w_{maxdestabilize}w_{destabilize}[DSR]}$$

$$\text{Flux of TNF translation} = k_{tl}[Tnfm]F_{repress\_tl}$$

sTNFR pre-treatment blocks TNFR signaling, as represented by a zero multiplier on TNFR activation.

$$\delta_{sTNFR} = \begin{cases} no \text{ sTNFR} \longrightarrow 0 \\ \text{sTNFR} \longrightarrow 1 \end{cases}$$

$$\text{Flux of TNFR activation} = (1 - \delta_{sTNFR})k_{activateTNFR}[TNFR][TNFpool]$$

BFA prevents secretion at the time of BFA treatment ($\tau_{BFA}$), as represented by a step-down multiplier on the flux of TNF secretion.

$$\delta_{BFA}(t, \tau_{BFA}) = 1 - H(t - \tau_{BFA})$$

$$\text{Flux of TNF secretion by a cell} = k_{secretion}\delta_{BFA}[TNF]$$

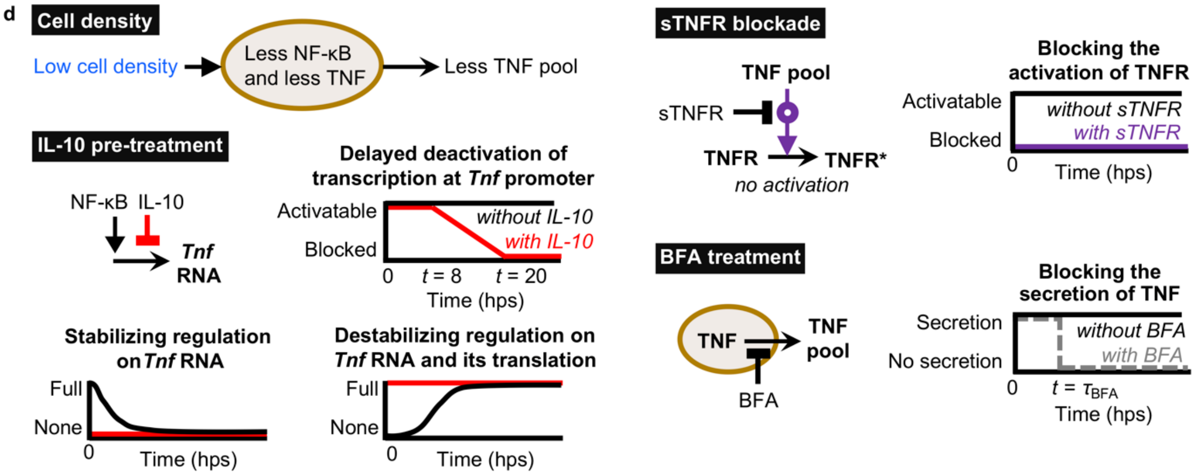


**Figure S3, continued.** (**d**) The diagrams summarize how perturbations are represented in the model. Low cell density leads to a decrease in LPS-induced TNF secretion for individual cells. IL-10 pre-treatment affects TNF expression by inhibiting sustained activation at the *Tnf* promoter post-LPS and by decreasing *Tnf* mRNA half-life through removing stabilizing regulation and promoting destabilizing regulation. sTNFR pre-treatment blocks TNF paracrine signaling. BFA treatment prevents TNF secretion, leading to intracellular TNF accumulation.

**State variables, parameters, and equations:**

**Supplementary Tables S2**, **S3**[1,4,16-19], **S4**, and **S5** are provided in accompanying spreadsheets.

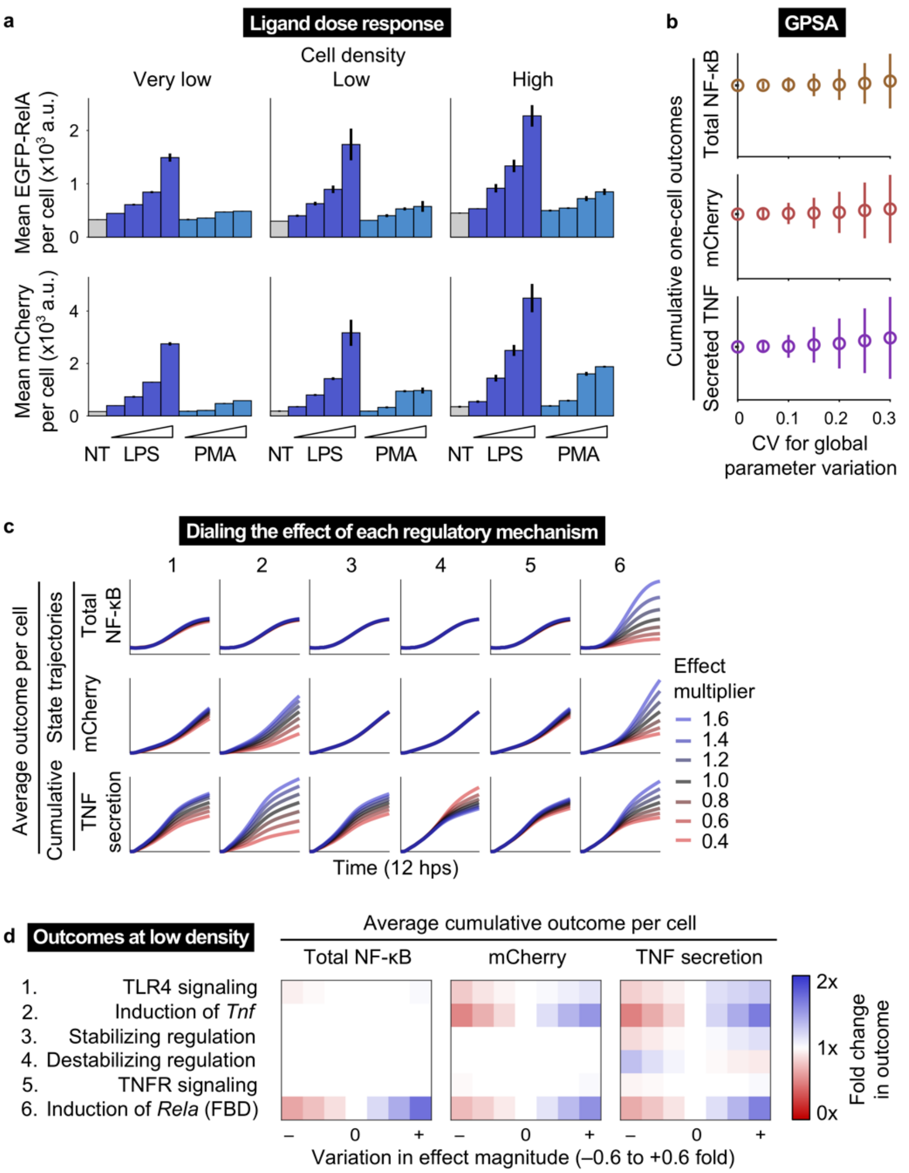


**Figure S4 | Model analysis.** (**a**) Flow cytometry data from **Fig. 4a**, shown here as bar graphs, for reporter proteins at 12 h post-LPS or PMA. Lines for each bar indicate +/– one standard deviation from the mean of three biological replicates. Ligand doses are 0.1, 1, 10, and 100 ng/ml. (**b**) Global parameter sensitivity analysis (GPSA). Values for the twelve estimated free parameters were varied simultaneously by randomly sampling from Gaussian distributions centered on the true-estimated values, with increasing coefficients of variation (CV; x-axis). One-cell simulations (*i.e.*, homogeneous case) were run at high density. Three metrics were used to summarize the outcomes, by integrating over 12 hps: total NF-κB, mCherry, and secreted TNF (flux). Y-axes are in linearly scaled a.u. specific to each plot, and are normalized to the base case values (at CV = 0). Line indicate +/– one standard deviation from the mean of 1,000 random samples per CV. Using the metrics shown, these outcomes can be interpreted as robust to global parameter variation within ~0.1–0.2 CV. (**c**) Outcomes from varying the effect magnitude (color-coded) for each mechanism in **Fig. 4b**. Population-mean reporter trajectories and cumulative secreted TNF are shown for a heterogeneous population at high density up to 12 hps. Cumulative secreted TNF is the total TNF secreted by a given time point, as determined by integrating the flux of secretion over time. Y-axes are in linearly scaled a.u. and are specific to each row of plots. (**d**) Analogous analysis to **Fig. 4b** for a population at low density. Trends are similar to those observed for a high density population.

### III. References


1 Cheng, Z., Taylor, B., Ourthiague, D. R. & Hoffmann, A. Distinct single-cell signaling characteristics are conferred by the MyD88 and TRIF pathways during TLR4 activation. *Sci Signal* **8**, ra69 (2015).

2 Basak, S., Behar, M. & Hoffmann, A. Lessons from mathematically modeling the NF-κB pathway. *Immunol Rev* **246**, 221–238 (2012).

3 Keogh, B. & Parker, A. E. Toll-like receptors as targets for immune disorders. *Trends Pharm Sci* **32**, 435–443 (2011).

4 Sung, M.-H. *et al.* Switching of the relative dominance between feedback mechanisms in lipopolysaccharide-induced NF- κB signaling. *Sci Signal* **7**, ra6 (2014).

5 Carpenter, S., Ricci, E. P., Mercier, B. C., Moore, M. J. & Fitzgerland, K. A. Post-transcriptional regulation of gene expression in innate immunity. *Nat Rev Immunol* **14**, 361–376 (2014).

6 Gaba, A. *et al.* IL-10−mediated tristetraprolin induction is part of a feedback loop that controls macrophage STAT3 activation and cytokine production. *J Immunol* **189**, 2089–2093 (2012).

7 Gais, P. *et al.* TRIF signaling stimulates translation of TNF-α mRNA via prolonged activation of MK2. *J Immunol* **184**, 5842–5848 (2010).

8 Kafasla, P., Skliris, A. & Kontoyiannis, D. L. Post-transcriptional coordination of immunological responses by RNA-binding proteins. *Nat Immunol* **15**, 492–502 (2014).

9 Brooks, S. A. & Blackshear, P. J. Tristetraprolin (TTP): interactions with mRNA and proteins, and current thoughts on mechanisms of action. *Biochim Biophys Acta* **1829**, 666–679 (2013).

10 Bonizzi, G. & Karin, M. The two NF-κB activation pathways and their role in innate and adaptive immunity. *Trends Immunol* **25**, 280–288 (2004).

11 Bode, J. G., Ehlting, C. & Häussinger, D. The macrophage response towards LPS and its control through the p38MAPK–STAT3 axis. *Cell Signal* **24**, 1185–1194 (2012).

12 Rajasingh, J. *et al.* IL-10-induced TNF-alpha mRNA destabilization is mediated via IL-10 suppression of p38 MAP kinase activation and inhibition of HuR expression. *FASEB J* **20**, E1393–E1403 (2006).

13 Parameswaran, N. & Patial, S. Tumor necrosis factor-α signaling in macrophages. *Crit Rev Eukaryot Gene Expr* **20**, 87–103 (2010).

14 Covert, M. W., Leung, T. H., Gaston, J. E. & Baltimore, D. Achieving stability of lipopolysaccharide-induced NF-κB activation. *Science* **309**, 1854–1857 (2005).

15 Sobol, I. M. Uniformly distributed sequences with an additional uniform property. *USSR Comp Math Math+* **16**, 1332–1337 (1976).

16 Werner, S. L., Barken, D. & Hoffmann, A. Stimulus specificity of gene expression programs determined by temporal control of IKK activity. *Science* **309**, 1857–1861 (2005).

17 Caldwell, A. B., Cheng, Z., Vargas, J. D., Birnbaum, H. A. & Hoffmann, A. Network dynamics determine the autocrine and paracrine signaling functions of TNF. *Genes Dev* **28**, 2120–2133 (2014).

18 Kalita, M. K. *et al.* Sources of cell-to-cell variability in canonical Nuclear Factor-κB (NF-κB) signaling pathway inferred from single cell dynamic images. *J Biol Chem* **286**, 37741–37757 (2011).

19 Maiti, S., Dai, W., Alaniz, R. C., Hahn, J. & Jayaraman, A. Mathematical modeling of pro- and anti-inflammatory signaling in macrophages. *Processes* **3**, 1–18 (2015).
